# Human library of cardiac promoters and enhancers

**DOI:** 10.1101/2020.06.14.150904

**Authors:** Ruslan M. Deviatiiarov, Anna Gams, Roman Syunyaev, Tatiana V. Tatarinova, Oleg Gusev, Igor R. Efimov

## Abstract

Genome regulatory elements play a critical role during cardiac development and maintenance of normal physiological homeostasis, and genome-wide association studies identified a large number of SNPs associated with cardiovascular diseases localized in intergenic zones. We used cap analysis of gene expression (CAGE) to identify transcription start sites (TSS) with one nucleotide resolution that effectively maps genome regulatory elements in a representative collection of human heart tissues. Here we present a comprehensive and fully annotated CAGE atlas of human promoters and enhancers from four chambers of the non-diseased human donor hearts, including both atria and ventricles. We have identified 10,528 novel regulatory elements, where 2,750 are classified as TSS and 4,258 novel enhancers, which were validated with ChIP-seq libraries and motif enrichment analysis. We found that heart-region specific expression patterns are primarily based on the alternative promoter and specific enhancer activity. Our study significantly increased evidence of the association of regulatory elements-located variants with heart morphology and pathologies. The precise location of cardiac disease-related SNPs within the regulatory regions and their correlation with a specific cell type offers a new understanding of genetic heart diseases.

## Introduction

The human heart has a sophisticated anatomical structure composed of different cell types that developed during embryogenesis or altered during aging or disease pathogenesis ^1,2^. All these features are orchestrated by precise Spatiotemporal regulation of gene expression by engaging numerous transcription factors (TF) at different time and loci, including promoter and enhancer regions ^3,4^. Evidence of high complexity of regulatory elements, significantly exceeding the number of genes in the human genome, leads to the widely accepted concept that mutations causing or related to diseases would be associated mainly with noncoding regulatory elements in the genome ^3,5^, in contrast to disease-causing mutations in protein-coding regions as in channelopathies ^6^. In cardiovascular diseases, this assumption was recently further supported by a combination of ENCODE data with Hi-C, strongly suggesting that causative mutations would be associated with cis-regulatory regions ^7–9^. Since cardiovascular diseases are the leading cause of morbidity and mortality in the United States and worldwide, we aimed to identify genomic localization of human heart-specific regulatory regions, hoping that such knowledge would facilitate the development of novel diagnostic and therapeutic strategies of heart disease.

Cap analysis of gene expression (CAGE) is a high throughput transcriptomic approach for transcription start sites (TSS) profiling at one bp resolution was used in FANTOM5 and ENCODE projects for accurate 5’ ends annotation in human, mouse, and other vertebrate genomes ^4^. Deep sequencing of CAGE libraries allows precise estimation of expression at the promoter level for coding and noncoding transcripts. This approach is useful for alternative promoter identification, which might affect final protein structure, or even estimate short TSS shifts linked to regulatory network features ^10,11^. Such cis-regulatory elements interact with enhancers and result in cell-type-specific expression profiles, while understanding of underlying molecular mechanisms and the network itself remains incomplete. It was shown that the bidirectional CAGE signals define positions of transcribed enhancers. Based on DNase I hypersensitive sites or histone modification signals, the transcribed enhancers usually function at a much higher frequency than the non-transcribed ones. These transcribed enhancers accumulate causative mutations as well and could be linked with CAGE promoters uncovering a comprehensive regulatory network^3^.

In this report, we present the results of deep CAGE analysis applied to 14 non-diseased donor human hearts. We demonstrate the existence of a large number of the additional TSS and enhancers specific to the human heart. The purpose of the project was to create a comprehensive annotated library of promoters and enhancers that are found in the human heart. This library contains four cardiac chambers: left and right atria and ventricles. Using this library, we identified a Genome-wide association study (GWAS) single nucleotide polymorphisms or SNP for leading cardiac diseases that are found in the regulatory regions. Additionally, comparing this database with specific cell type expression in FANTOM5 data allowed us to identify cardiac cell-type-specific promoters, which can be used as novel markers for cell identification or therapeutic targets.

## Results

### Human heart CAGE library identified novel promoters and enhancers

We collected tissue samples from non-diseased human donor hearts (**Supplementary Table 1**) from four cardiac chambers: left and right, atria, and ventricles. We used a total of 41 samples from 14 hearts for CAGE library construction, sequencing, mapping to genome, and TSS identification. Based on this CAGE data, we identified 49,179 new heart CAGE robust decomposition peak identification (DPI) clusters, which had a 78.6% overlap with FANTOM5 annotation, and the remaining 10k were new previously unknown peaks (**Fig. 1a**). These peaks were classified into TSS-like according to nucleotide distribution against the Eukaryotic Promoter Database (EPD) (29,598 TSS). They resulted in 27,295 TSS-like clusters in heart, where 2,750 had no overlap with FANTOM5. Bidirectionally expressed CAGE TSS (CTSS) were used for cardiac-active enhancer annotation (n=6,147) and compared to FANTOM5 (**Fig. 1a**). These heart CAGE based regulatory elements had features of active promoters and enhancers (**Fig. 1b-d**; **Supplementary Fig. 1**) and were used in all remaining analyses. Notably, heart CAGE peaks accumulate DNase I hypersensitive sites, and histone modification marks signals at a higher level than FANTOM5 peaks, while heart ATAC-seq signal enrichment was expected and representative in case of enhancers (**Supplementary Fig. 2**). These observations, together with high conservation scores and CpG overlap proportion, suggested the housekeeping nature of defined heart CAGE clusters. Almost all peaks classified as TSS were associated with transcripts and related genes, while in the case of DPI clusters and enhancers, 82% and 20% could be linked respectively (**Fig. 1e**).

**Fig. 1.**
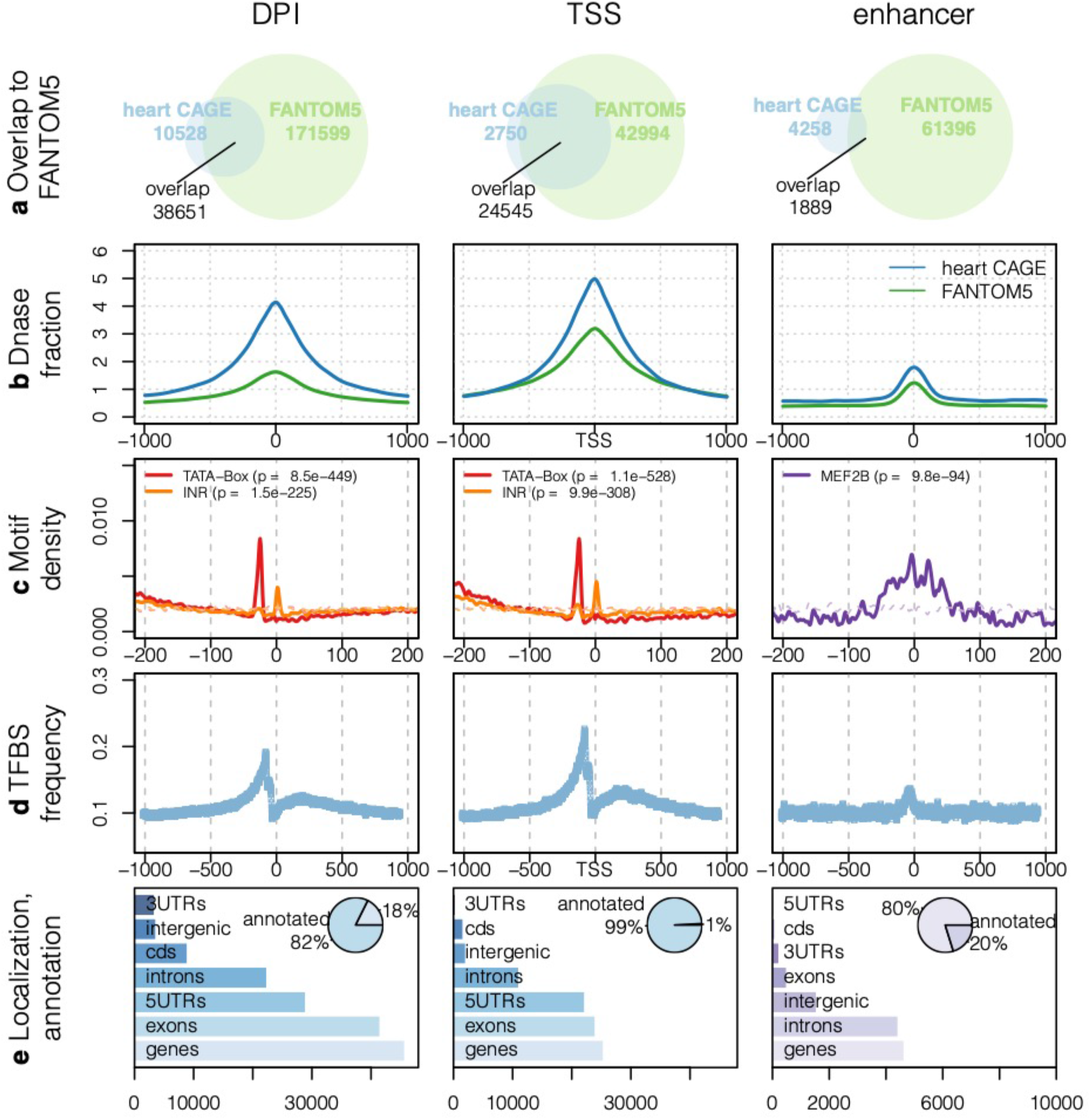
Discovery of new promoters and enhancers in the human heart. **a**, Heart CAGE library of promoters (robust DPI and TSS-like DPIs) and enhancers overlap with FANTOM5 clusters, but significant numbers are heart CAGE specific. Notably, TSS-like overlap is the highest, while in the case of enhancers it is low, representing the conservation level of these elements. **b**, Dnase cluster density shows open chromatin transcription. Heart promoters and enhancers have similar trends to FANTOM5 data but accumulate signals at higher levels. **c**, Location breakdown of Pol2 motifs within discovered heart elements: TATA-box and INR are shown. In the case of enhancers MEF2B (Myocyte-specific enhancer factor 2B) was the top enriched motif. **d**, Transcription factor binding sites (TFBS) frequency in heart CAGE library. Discovered TFBS distribution is a sign of promoter and enhancer function. **e**, Annotation of heart library elements by the association with gene models. Most of the heart CAGE peaks were connected to Gencode v33 genes, but remaining peaks show potential for annotation improvement.

Since there was a significant overlap of heart CAGE peaks with FANTOM5, we assigned available ontologies (cell line, disease, organ) to robust DPIs. Interestingly, enrichment results showed embryo related features in defined heart CAGE peaks (**Supplementary Fig. 3**); cardiovascular system-related tags were enriched as well. Disease and Cell ontology enrichment results were consistently reporting “cancer” and “embryonic cell” as the top hits, also including heart-related diseases, myoblasts, fibroblasts, and other expected cell types. These results agree with the heart composition profile since we found mesodermal cells, muscle stem cells, smooth muscle cells, electrically active/responsive cells, fibroblasts, and other cell type-specific promoters in the heart (**Supplementary Fig. 3**).

The total number of genes active in the heart was 13,849 (**Supplementary Table 2**), resulting in 3.6 robust DPI to gene ratio and 2.47 TSS per gene. Only 30.0% (4,149) genes have a single CAGE peak (**Fig. 2a**), but a majority of genes (70%) had multiple DPI and TSS. Moreover, while in general heart CTSS distance to available transcripts models was within 98 bp, numerous outlier peaks could be found (n=6,003, **Fig. 2a**). Genes with such “distal” CAGE peaks were associated with numerous heart-related diseases including cardiomyopathy, hypertrophic cardiomyopathy, cardiovascular system disease (**Fig. 2b**), and related to regulation activity, like transcription factor binding or co-regulator activity (**Fig. 2c**).

**Fig. 2.**
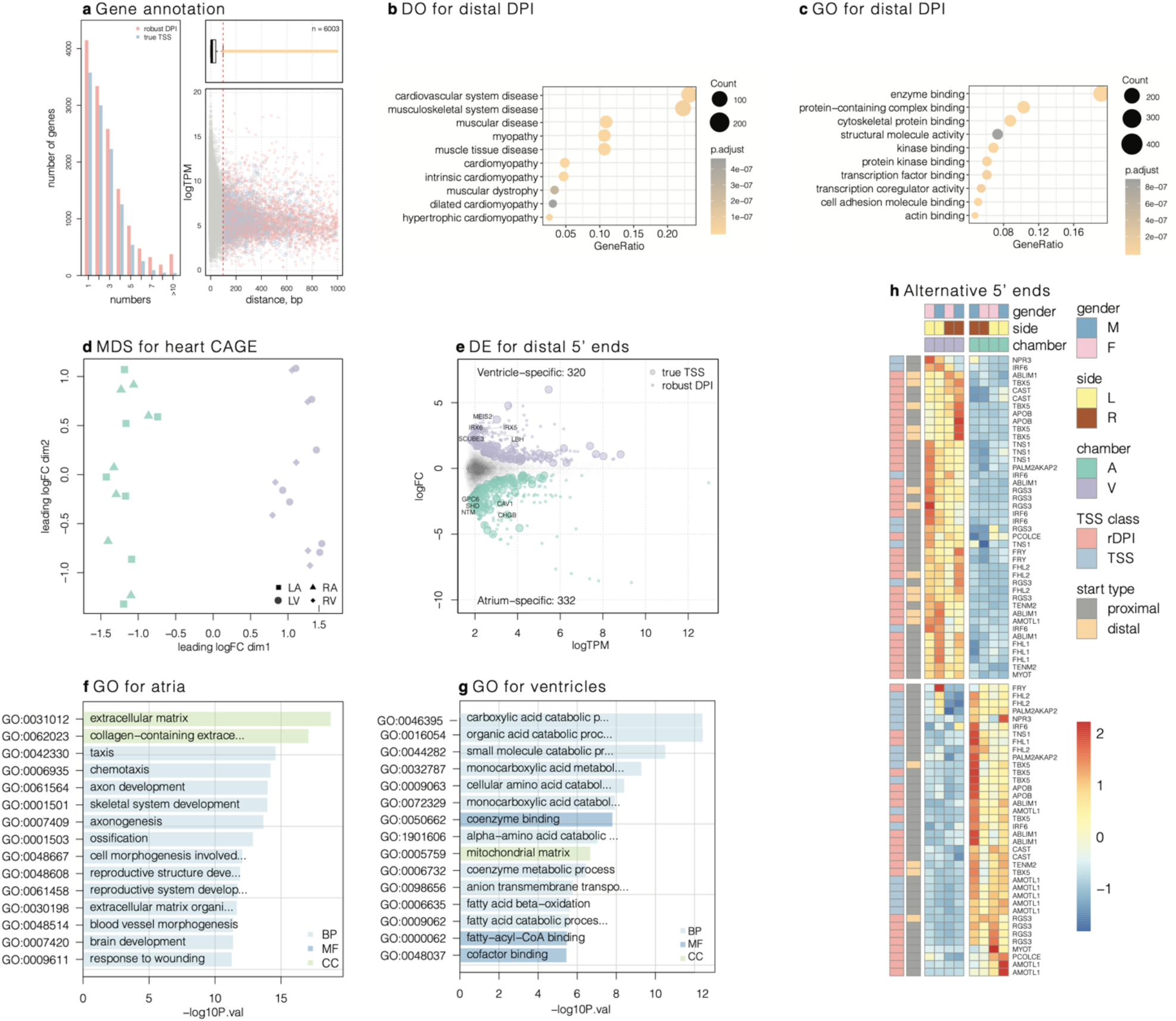
Promoter level expression atlas in heart chambers defines specific TSS. **a**, Gene annotation by CAGE peaks. Within 13850 unique genes, 9701 have more than one DPI and 7472 – more than one TSS-like peak. Distances of 6003 CAGE peaks to related transcripts longer than 98 bp (“distal” peaks), where 2060 are TSS-like. **b-c**, Genes with distal CAGE peaks improved 5’ ends have significant importance in heart diseases and play regulatory roles through protein binding. **d**, MDS plot for heart CAGE clusters based on young samples (<40 y.o.). Ventricular and atrial libraries are clearly separated. **e-f**, Functional annotation of Gene Ontology (GO) of chamber specific genes (n=583 in the ventricle, and 985 in the atrium). BP: biological process, MF: molecular function, CC: cellular component. **g**, Differentially expressed “distal” CAGE peaks between atrium and ventricle (robust DPI and TSS-like), the top five specific genes are shown. **h**, Promoter level heatmap for genes with atrial and ventricular specific expression (|logFC|>1, FDR<0.05). Repeating gene names signify the presence of multiple promoters. The legend shows scaled TPM values distribution.

### CAGE library identified cardiac-specific alternative promoters

CAGE, as a transcriptomic approach, allowed estimating expression precisely at the promoter level and resulted in a clear separation of ventricular and atrial samples in our study (**Fig. 2d-e**). In total, we obtained 2,111 atria and 1,589 ventricle specific CAGE peaks, where 830 and 491 are TSS-like, respectively (|logFC|>1, FDR<0.05; **Supplementary Table 2**). Functional analysis of specific genes revealed neural system related ontology in atria while catabolic, and oxidation processes were enriched in the ventricles (**Fig. 2f-g**). FANTOM5 ontologies-based enrichment for specific groups resulted in embryo type tags for atria and heart tags (primary circulatory organ, cardiac valve, endocardium, etc.) in ventricles. In contrast, heart and vascular diseases were predictably overrepresented in both chambers (**Supplementary Fig. 4**). Fibroblasts, myoblasts, multi-potent muscle stem cells, electrically active/responsive, and muscle precursor cells were enriched in the atria, while epithelial and fat cells in ventricles (**Supplementary Fig. 4**). Genes annotated by heart CAGE peaks and specifically expressed in chambers had cardiovascular system disease and cardiomyopathy as top tags in atria and ventricles, respectively (**Supplementary Fig. 5**). These genes had clear functional separation at pathway level through KEGG annotation: atrial genes participate in PI3K–Akt, TGFβ, AGE–RAGE signaling; ventricular genes are involved mostly in carbon metabolic pathways, including PPAR and fatty acid degradation (**Supplementary Fig. 5**). Motif analysis for specific promoter regions resulted in androgen receptor (AR), NFIX, ZNF263 top transcription factors in the atria, and Esrrb, ESRRA, NR4A2, HNF4A, ESR2 in ventricles (**Supplementary Fig. 5**).

We also found that hundreds of newly annotated heart CAGE peaks could be classified as “distal” and specifically expressed in ventricles and atria: 320 and 332, respectively (**Fig. 2e**). There are 13 genes, which have more than one CAGE peak with the opposite specificity (**Fig. 2h**), and at the same time, some of them are characterized as TSS-like or “distal.” Some representative examples are illustrated in **Supplementary Fig. 6-7.** The utilization of alternative promoters allows alternative splicing resulting in multiple transcript variants encoding different protein isoforms.

One of the critical regulators of heart development, TBX5, had one CAGE peak classified as TSS-like, which was located close to the existing gene model and expressed specifically in the atria. Still, alternative downstream CAGE peak had ventricle-specific expression (**Supplementary Fig. 6**). TBX5 acts as a key transcription factor during embryonic development, and its mutations have been linked to several cardiac defects and diseases, including atrial fibrillation and cardiac septal defect ^12,13^. PITX2 gene encodes a regulator of TBX5 ^14^ and has also been implicated in the pathogenesis of atrial fibrillation and cardiomyopathy ^15^. Interestingly, PITX2 has multiple CAGE peaks with left atrial specificity (**Supplementary Fig. 6)**

Another intriguing example, KALRN gene encodes several isoforms by using alternative transcription start and termination sites (**Supplementary Fig. 6, 14, Table 6-7**).^16,17^ CAGE sequencing analysis confirmed the heart activity of at least two of these alternative starts of transcription, 552,218 bp apart. The two start sites were located in positions 124,032,456 (Site A) and 124,584,674 (Site V); Based on CAGE, Site A is approximately equally active in both atria and ventricles (15.66:12.15), while Site V is almost 3-fold more active in the ventricles (8.94:25.13). Site V corresponded to the shorter isoform (V), encoding the 1288 aa long protein, while Site A corresponded to the longer isoform (A), encoding the 2988 aa long protein. The additional 2,000 amino acid long sequence contains five copies of the Spectrin repeat, Sec14p-like lipid-binding domain, and an extra set of RhoGEF/PH2_Kalirin_Trio/SH3, already included in the shorter isoform V. Spectrin repeats of the isoform A are used for interaction of KALRN with several proteins, including inducible nitric oxide synthase.^18–20^ Nitric oxide mediates multiple physiological and pathophysiological processes in the cardiovascular system, especially metabolism^21^, its crucial role in the development of cardiovascular diseases has been demonstrated by numerous human and animal studies.^22–24^ The isoform A contains an intronic SNP, rs9289231, associated with early-onset coronary artery disease (CAD).^25^ It has been proposed that the role of KALRN in CAD susceptibility is through its interaction with the inducible nitric oxide synthase.^25^

Calpastatin or CAST is a calpain (calcium-dependent cysteine protease) inhibitor. Calpain family has been shown to play an essential role in the pathogenesis of hypertrophy; for example, calpain inhibition has been shown to modulate the process of ventricular hypertrophy. CAGE data indicate alternative splicing domains between atria and ventricles, which might explain why inhibition by calpastatin is more effective in the ventricles than atria ^26^ (**Supplementary Fig. 6**).

Gene IRF6 has two transcription initiation regions: 2 and 5 DPI are atria and ventricle specific, respectively (**Supplementary Fig. 6**). In these two cases, the final protein product remains unchanged, but in the case of ABLIM1, AMOTL1, CAST, RGS3, the structure is likely to be changed (**Supplementary Fig. 6**). IRF6, ABLIM1, and AMOTL1 are involved in the developmental process of the heart. All these examples could be further investigated through the Zenbu page (https://fantom.gsc.riken.jp/zenbu/reports/#Human_Heart_CAGE_A).

### SNP related to cardiac diseases accumulate in the TSS region

The intersection of heart CAGE clusters with heart-related GWAS SNPs resulted in the accumulation of variants in DPI regions. There are 1.7-85.4 % (9.86% in median) of SNPs on primary chromosomes located in heart promoter regions defined in this study, 1.4-77.1% in “robust” DPI, as defined by FANTOM5 pipeline (**Fig. 3a**). After normalization by the total length of the heart, CAGE DPI regions, and Gencode v33 coding sequence (CDS) showed the highest proportion of heart GWAS SNPs. We also found SNPs distribution peak close to TSS defined by heart CAGE (**Fig. 3b**), as well as for several heart diseases (**Fig. 3c-g**). In total, 4,302 and 12 unique heart GWAS SNPs were identified in heart CAGE DPI and enhancers, respectively. In DPI regions (±250 bp), the average MAF is 0.027, compared to 0.117 genome-wide except DPI. Distribution of MAF and SNPs density near the TSS-like areas is shown in **Supplementary Fig. 8**.

**Fig. 3.**
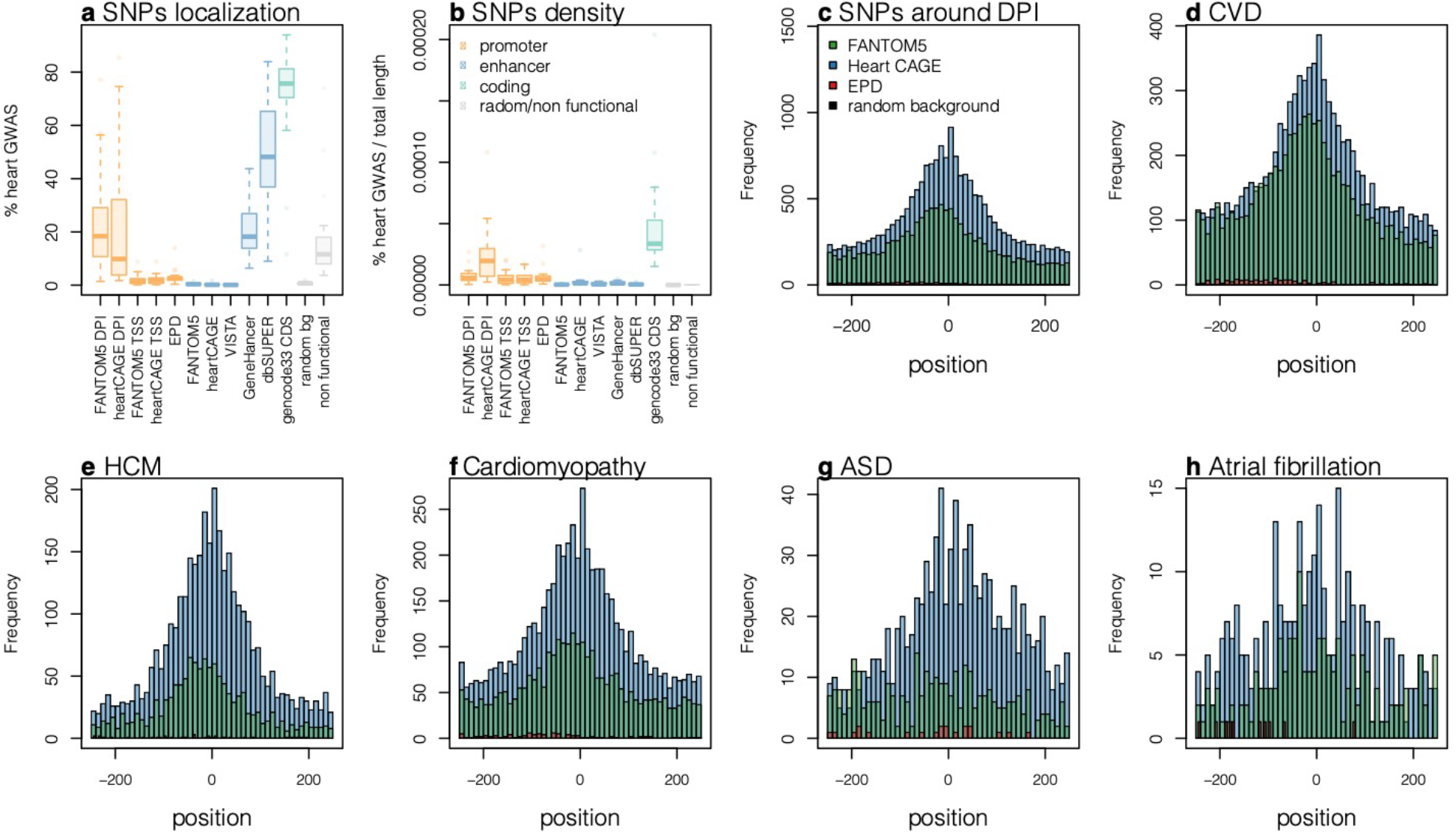
Heart CAGE promoters accumulate disease causative variants. **a**, Genomic localization of heart GWAS SNPs for primary chromosomes. Random bg - random background clusters n=30k, l=400bp. **b**, Distribution of heart GWAS SNP frequency in heart CAGE defined TSS regions based on DPI regions. **c-h**, Examples of SNP distribution for specific heart diseases show their clustering near TSS. CVD: Cardiovascular disease, HCM: Hypertrophic cardiomyopathy, ASD: Atrial septal defect.

### Quality of annotation allowed the identification of novel regulatory elements

The prediction quality of promoters and enhancers was assessed from the analysis of nucleotide consensus, DNA methylation, and characteristic motifs in the vicinity of the TSS (**Fig. 4**). To compare the two groups (promoters and enhancers), we selected two subsets of data. To represent the promoters, we have chosen those TSS that were annotated as “true,” then we have selected the start site with the highest CAGE expression as the “best” TSS per gene (10,206 TSS). The distribution of frequencies of nucleotides (**Fig. 4a**) is consistent with the well-defined start of transcription. There are peaks of A and T at TSS-30 and pronounced G-richness of the transcribed region; all of these observations were well-documented in prior studies and across multiple organisms ^27–32^. Enhancer positions were predicted all over the genome, with 83% of them located inside a transcribed sequence. We have excluded those enhancers that overlapped with exons, 5’ UTR, 3’ UTR, and promoters. Ten thousand eight hundred thirty-two enhancer-TSS sequences satisfy these conditions. Statistical patterns of nucleotides around the “enhancer-TSS” (**Fig. 4b**) differed from a typical TSS distribution showing narrow and pronounced peaks at the TSS and TSS-30. DNA sequence around the enhancer-TSS showed a GC-rich 500 nucleotide long “bubble.”

**Fig. 4.**
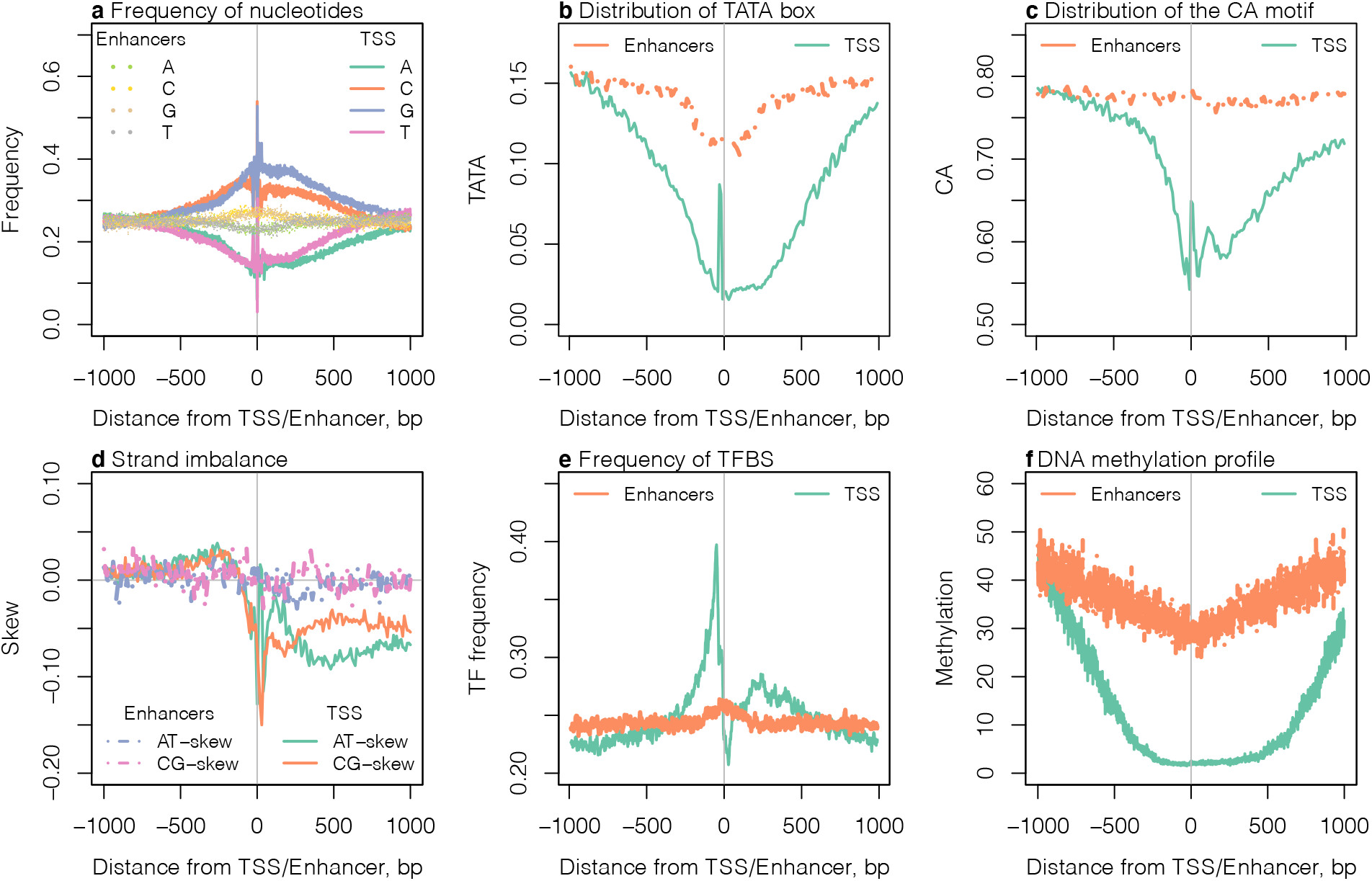
Comparison of genomic regions in the vicinity of TSS and enhancers. **a**, Frequency of nucleotides. **b**, Strand imbalance, measured as AT and CG skews, averaged in 10 nucleotide windows. **c**, Distribution of TATA box. **d**, Distribution of the CA motif, the initiator consensus signal. **e**, Frequency of TF binding sites. **f**, DNA methylation profile measured as Beta (mean).

TATA-box is usually located at −30 upstream from the TSS and is particularly important. CAGE-annotated promoters showed a peak of TATA-like sequences (“TATA”, “TAAA”, and “ATAT”) at the window [TSS-40, TSS-20] (**Fig. 4b**). Enhancers had a dip in the frequency of TATA at TSS. Another remarkable feature of the TSS is the CA dinucleotide motif located at the start of transcription (**Fig. 4c**). CAGE-predicted TSS also had an elevated frequency of this motif, while enhancers lacked this trend. Compositional strand asymmetry is a characteristic feature of important genomic regions, such as starts of transcription, tRNA insertion sites, and transposable elements ^33^. Peaks in CG-skew and AT-skew 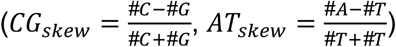 were previously associated with TSS ^34,35^. At the human TSS, there was a narrow peak of AT-skew and similarly small dip of CG-skew (**Fig. 4d**). The trends in enhancers were different and much more subtle: they had small approximately symmetrical peaks of AT-skew and CG-skew ~300-200 nucleotides upstream/downstream of the enhancer-TSS (Fig. 4D)

Another remarkable feature of promoters - the peak of the density of transcription factor binding sites (TBFS) upstream of the start of transcription (**Fig. 4e**). This peak corresponded to the core promoter, typically found at [TSS-200, TSS]. TFBS distribution (**Fig. 4e**) had a sharp spike in the core promoter region [TSS-200, TSS], followed by a drop in the 5’ UTR, and then there was a second smaller peak at TSS+300 nucleotides. As with all other features, the trends near the enhancer-TSS were much less pronounced: only a slight increase of TBFS density at enhancer-TSS. A natural question if the sequence motifs near TSS and enhancer-TSS are the same? What motifs are associated with elevated levels of transcription? We have conducted an analysis of sequence motifs using Match ^36^ for known TFBS and cisExpress^27,30^ for de-novo discovery. For both TSS and enhancer-TSS, the most common and essential for transcription motifs were made of cytosines and guanines: GGGGC, GGGGG, GGCGG (and their reverse complements) ATAAT (and variations).

Trends of the methylation profiles could also be used to validate the quality of TSS prediction ^28,37^. There was a drop in the DNA methylation that starts immediately upstream of the TSS. Since the gene bodies are generally more methylated,^38^ the resulting pattern is U-shaped (**Fig. 4f**)

## Discussion

We aimed to create a comprehensive map of a human promoterome, which allows grasping the general topology of regulatory landscape of the genome and focusing on its specific players and features. This map will allow us to detect transcriptional changes in many cardiac physiological states and different stages of disease pathogenesis. It will also help develop novel interventions and therapies, targeting a cardiovascular network of regulatory elements of the genome. Additionally, the expression of regulatory elements of four cardiac chambers can be a useful tool to determine the maturity of the induced pluripotent stem cells that are differentiated into atrial or ventricular phenotype in general or more subtle phenotypes of sinoatrial nodal cells, Purkinje cells, etc.

The library of human cardiac promoters and enhancers presented here contains over forty thousand regulatory elements that control transcription of thirteen thousand genes expressed in the heart. Our results showed a substantial amount of cardiac disease-associated genetic variants were located in noncoding regulatory regions (**Fig. 3a**),^39^ rather than in protein-coding regions. As our GWAS SNP distribution showed, several variants associated with the disease were also enriched in the promoter region of the gene. This enrichment supports the importance of studying the regulatory elements and their association with disease. It also presents a novel concept of heritable cardiovascular diseases, different from the traditional idea of disease-causing mutations within the protein-coding regions (such as channelopathies, when mutations lead to electrogenic ion channel gain or loss of function and life-threatening arrhythmias). Our findings show that arrhythmias, heart failure, and developmental disorders could be caused by mutations in the regulatory elements.

Identifying heart-specific transcription factors and transcription factor binding sites allows designing new therapeutic targets that target the regulation of genes that lead to cardiac diseases. For example, transcription factors are involved in stress regulation of the adult heart that can result in hypertrophy, fibrosis, or ischemic injury ^40,41^. Since the majority of the mechanisms work in opposition, an imbalance of the network of regulatory factors can lead to the pathogenesis of a disease. Rebalancing the network of transcriptional control by targeting TFBS could serve as a novel strategy of treatment of cardiovascular diseases ^42^.

### Future Directions and Limitations

Heart muscle comprises many different cell types, including myocytes, fibroblasts, endothelial cells, adipocytes, macrophages, neurons, and many others. The current tissue level CAGE approach provides limited insight into the cell-type-specific activity of promoters and enhancers. To address this limitation and to achieve a higher resolution of our library, implementing the single-cell CAGE is highly desirable. This analysis would enable us to map the transcriptome of the entire heart, including the conduction system, vascular system, neural and immune systems, and electrical system at a single-cell resolution ^43^. Additionally, it would aid in linking the physiological and electrical properties of cardiac cells to their specific regulatory element signature and novel molecular markers.

The ultimate goal is to create a comprehensive catalog of functionally supported human regulatory elements that are well annotated in terms of the cell types, active regulatory elements, the genes that each element regulates, and the degree of activation of each element.^44^

## Online Methods

### Donor Heart Procurement and Demographic

Adult human cardiac tissue for this study was procured from the Washington Regional Transplant Community (Falls Church, VA) as de-identified donor human hearts, which were not accepted for transplantation. The cause of death was non-cardiogenic. All protocols were approved by the George Washington University Institutional Review Board. Explanted hearts were arrested at the time of procurement with the cardioplegic solution and cooled down to +4 °C for transportation to the research laboratory, which was typically less than 2 hours.

Demographic breakdown of the procured hearts was as follows: donor hearts less than 40 years of age were included in the young category that contained three males and four females (**Supplementary Table 1**). Donor hearts older than 40 years were included in the old category with four males and three females. In total, we have sequenced 41 samples from 14 hearts. The “old” group (>40 y.o.) was sequenced first. The sequencing was performed on the nAnTi-CAGE platform, and tissues from the left atrium and ventricle were collected and sequenced. After seeing the difference between atria and ventricle, we decided to explore the difference between right and left side as well, by including all four chambers into the new sequencing as well as expand on the age range of our samples, thus sequencing the second group of “young” hearts (<40 y.o.). This set of samples was sequenced on a new platform, ssCAGE.

### Tissue Preparation and Sequencing

Upon delivery to the laboratory, small pieces, 0.5 cm^3^, were dissected from the left and right atrial free wall and the right and left ventricular base. The tissue was stored in RNAlater (Invitrogen) overnight at +4 °C and then stored at −80 °C until RNA extraction. Total mRNA was extracted from 40 mg of tissue with RNeasy Fibrous Tissue Mini Kit (Qiagen) according to the manufacturer’s protocol. RNA yield and purity were quality checked with Agilent Bioanalyzer and Qbit to satisfy RIN >7 and total RNA amount 3-5 μg. For the old group, total RNA was submitted for no-amplification non-tagging CAGE (nAnT-iCAGE) library preparation and sequenced on Illumina HiSeq2500 High-Throughput mode with 50nt single end and with 6M reads per sample ^45^. For the young group, total RNA was sequenced on Illumina HiSeq2500 by one-shot loading with 80nt and over 25 M reads per sample ^46^.

### CAGE Data Processing

Sequenced reads were checked for quality with FASTQC, then trimmed for quality and length with fastx_trimmer (-Q33) and adapters removed with trimmomatic ^47^. Reads mapped to human ribosomal RNA locus (U13369.1) were removed. Clean reads were mapped on hg38 primary chromosomes with Burrows-Wheeler Aligner ^48,^ and unmapped reads were remapped with HISAT2 ^49^. Mapped reads were converted into CTSS (CAGE TSS) using CAGEr Bioconductor package (sequencingQualityThreshold = 10, mappingQualityThreshold = 20, removeFirstG = TRUE, correctSystematicG = TRUE) to count 5’ end of the mapped CAGE reads at single base-pair resolution ^50^.

Due to the heterogeneous profile of TSS clusters, decomposition peak identification (DPI) that uses independent component analysis, DPI program ^4^ was used to define a set of reference peaks applying on all 41 samples. Robust DPIs (TPM>1) were then linked with genes through a custom-developed CAGE_peak_annotation package. Augustus, Genscan, NCBI RefSeq, and Gencode v33 models were downloaded from Gencode and UCSC Table Browser. TomeTools TSSClassifier ^4^ was used to classify robust DPI peaks and to define a set of TSS-like peaks. DPI and TSS peaks were further analyzed with edgeR for differential expression analysis and with topGO, clusterProfiler for GO terms. For differential expression data analysis, “young” samples (<40 y.o.) were used. The WGCNA package was applied for co-expression clusters identification on both experimental groups separately with soft threshold 9 and 14 for groups one and two, respectively.

Enhancers were called by FANTOM5 scripts ^3^ and linked to genes, as described by Andersson et al., 2014. Motif enrichment analysis was performed in MEME Suite ^51^ with core, vertebrates, and Pol2 JASPAR motifs collections 2018.

Signal data for epigenetic marks of Pol2, H3K27ac, H3K4me3, H3K4me1 was obtained from the ENCODE project. ATAC seq data for the human heart was obtained from Broad Institute’s Cardiovascular Disease Knowledge Portal ^52^. Conservation scores phastCons from 100-way alignment and CpG islands regions were accessed from UCSC for hg38. Fractions of signals were counted using bedtools v2.28.0. Vista enhancers ^53^, super-enhancers ^54^, and *H. sapiens* promoters EPD ^55^ were obtained from related sources. UCSC LiftOver tool applied in case if hg38 annotation was unavailable.

Heart GWAS data for hg38 assembly was obtained from Ensembl BioMart (**Supplementary Table 3**). NCBI dbSNP Build 151 data downloaded from UCSC. The frequency for heart GWAS SNP was counted within ±200 bp from newly defined or known TSS. Percentage of heart GWAS SNP overlapping different genomic regions (promoters, enhancers, protein-coding, nonfunctional) was counted for each chromosome separately.

All obtained heart CAGE data, annotations, and analysis results were integrated on the Zenbu-reports platform ZENBU 3.0.1. which is provided here as an open-source database. To compare our regulatory elements data to public SNP (**Supplementary Table 3**) and DNA methylation data we used several public gene expression datasets (https://www.ebi.ac.uk/arrayexpress/experiments/ 6814, 513, 4344, 3716, and 3358), and the iMETHYL database (http://imethyl.iwate-megabank.org/).

### Statistical Analysis

We analyzed regulatory sequence motifs using Match^36^ for known TFBS and cisExpress 27,30 for *de novo* discovery. We used the Match tool^36^, with matrix similarity at least 0.95 and core similarity 1, and calculated the distribution of density of TFBS. The clustering of the TBFS was conducted based on the positional frequency. First, we calculated position-specific density profiles for each TFBS in the TRANSFAC database and excluded the rare motifs (those that appear less than in 1/10 of the analyzed sequences). Then we computed Pearson’s correlation coefficients between densities of motifs in the window of 10 nucleotides. Then we used hierarchical clustering in R (hclust), using the Ward method and (1-correlation)/2 as a distance. To test the significance of differences between clusters of TFBS, we used the Kruskal-Wallis test (R Kruskal.test), and the Wilcoxon rank-sum test with continuity correction (R wilcox.test) to assess pairwise differences.

For the *de novo* motif discovery, we used the cisExpress^27,30^ algorithm. This approach is based on the calculation of the Z-score, showing the strength of the influence of having a nucleotide sequence *w* in the positional window *k*. The size of the window (*w*) is a parameter that is selected based on the quality of the genome annotation and properties of the organism and the experiment.

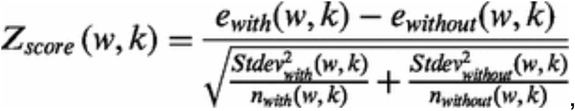

where *e*_*with*_*(w,k)* and *e*_*without*_*(w,k)* are average gene expression values with and without the word *w* in the position *k*; *Stdev*_*with*_*(w,k)* and *Stdev*_*without*_*(w,k)* are the standard deviations of gene expression values; *n*_*with*_*(w,k)* and *n*_*without*_*(w,k)* are the numbers of sequences of genes containing and not containing word *w* in the *k*th window. Words with *Z*-scores above a predefined threshold are stored as primary motifs. Groups of similar motifs discovered within one window are then merged, resulting in longer and/or more ambiguous motifs. This part of the algorithm produces the motif in the form of a consensus sequence and includes the position of the window where it was discovered.

### Data Availability

Human heart CAGE data is available on GEO, number GSE150736: raw reads, CAGE DPI, and enhancers are available (**Supplementary Table 2**). For interactive data browsing, we created a customized database using the Zenbu reports platform for TSS, enhancer, and currently available heart-related GWAS SNPs: https://fantom.gsc.riken.jp/zenbu/reports/#Human_Heart_CAGE_A.

User manual on how to navigate the human heart CAGE library is attached in supplementary files.

### Ethical Approval

This study was approved by the George Washington University Institutional Review Board.

## Supporting information

Supplement 5 Zenbu User Manual

## Abbreviations

A: atrium
ASD: atrial septal defect
bg: background
bp: base pair
CAD: coronary artery disease
CAGE: cap analysis of gene expression
CDS: coding sequence
CTSS: CAGE transcription start site
CVD: cardiovascular disease
DO: Disease Ontology
DOID: Disease Ontology ID
DPI: decomposition peak identification
EPD: eukaryotic promoter database
FANTOM5: functional annotation of the mammalian genome 5th version
GENECODE: genetic encyclopedia of DNA elements
GO: Gene Ontology
GWAS: genome-wide association study
HCM: hypertrophic cardiomyopathy
INR: initiator
KEGG: Kyoto Encyclopedia of Genes and Genomes
LA: left atrium
logFC: log2 fold change
LV: left ventricle
MAF: minor allele frequency
MDS: multidimensional scaling
SNP: single nucleotide polymorphism
TF: transcription factor
TFBS: transcription factor binding site
TIR: transcription initiation region
TSS: transcription start site
V: ventricle

## Links

Library of heart promoters and enhancers: https://fantom.gsc.riken.jp/zenbu/reports/#Human_Heart_CAGE_

DPI https://github.com/hkawaji/dpi1/

enhancers https://github.com/anderssonrobin/enhancers

annotation https://github.com/Deviatiiarov/CAGE_peak_annotation

TSS classification https://sourceforge.net/projects/tometools/

UCSC data http://hgdownload.soe.ucsc.edu/goldenPath/hg38/database/

FANTOM5 data https://fantom.gsc.riken.jp/5/datafiles/reprocessed/hg38_latest

## Acknowledgments

This study was funded by Leducq Foundation (project RHYTHM to IRE and AG), National Institutes of Health (3OT2OD023848, R01 HL126802, U01 HL141074, R44 HL139248 to IRE) and Russian Scientific Foundation grant 18-71-10058 (to RS), and University of La Verne Faculty Development fund (to TT).

## Author contributions

AG and IE collected samples; OG performed sequencing; RD and TT conducted data analysis; RD, AG, RS, TT, OG, and IE wrote the manuscript.

## Materials & Correspondence

Correspondence and material requests should be addressed to Oleg Gusev, Tatiana Tatarinova and Igor Efimov.

