## Supplement 1 for "Human library of cardiac promoters and enhancers"

### Supplement for:
### Human library of cardiac promoters and enhancers

Ruslan M. Deviatiiarov[1], Anna Gams[2], Roman Syunyaev[3,2,10], Tatiana V. Tatarinova[4-7*], Oleg Gusev[8*], Igor R. Efimov[2,3,9*]

[1] Institute of Fundamental Medicine and Biology, Kazan Federal University, Kazan, Russia
[2] The George Washington University, Washington, DC, USA
[3] Moscow Institute of Physics and Technology, Dolgoprudny, Russia
[4] Department of Biology, University of La Verne, La Verne, California, USA
[5] IPPI RAN, Moscow, Russia
[6] Institute for General Genetics, Moscow, Russia
[7] Siberian Federal University, Krasnoyarsk, Russia
[8] Division of Genomic Technologies, Center for Life Science Technologies (CLST), RIKEN, Yokohama, Japan
[9] IHU LIRYC, Site Xavier Arnozan, Avenue du Haut Lévêque, Pessac, France.
[10] Sechenov University, Moscow, Russia

*NOTE: **Supplementary Table 2 & 3** - separate .txt files

**Supplementary Fig. 6** - separate .pdf file

### Sample description

**Supplementary Table 1. Sample Demographics.** Human donor hearts were broken down into two groups: young (<40 y.o.) **(upper table)** and Old (>40 y.o.) **(lower table)**.

**Young (<40 y.o.)**

| Sex | Age | Body Mass Index | Left Ventricular Ejection Fraction (%) | Cause of Death | Cardiac Chamber |
|---|---|---|---|---|---|
| Male | 26 | 26.4 | 65 | Anoxia/Drug Intoxication | LA, RA, LV, RV |
| Male | 28 | 18.3 | 25-30 | Anoxia/Cardiovascular | LA, RA, LV, RV |
| Male | 34 | 38.2 | 65 | Anoxia | LA, RA, LV, RV |
| Female | 26 | 36.1 | 60-65 | Infectious Disease | LA, RA, LV, RV |
| Female | 35 | 31.5 | 65 | Cerebrovascular Accident/Stroke | LA, RA, LV, RV |
| Female | 36 | 31.7 | N/A | Cerebrovascular Accident/Stroke | LA, RA, LV, RV |
| Female | 39 | 36 | 45-50 | Anoxia | LA, RA, LV, RV |

**Old (>40 y.o.)**

| Sex | Age | Body Mass Index | Left Ventricular Ejection Fraction (%) | Cause of Death | Cardiac Chamber |
|---|---|---|---|---|---|
| Male | 43 | 27.8 | N/A | Head Trauma/Blunt Injury | LA, LV |
| Male | 51 | 26.3 | N/A | Cerebrovascular Accident/Intracerebral Hemorrhage/Stroke | LA |
| Male | 60 | 22.8 | N/A | Cerebrovascular Accident/Intracerebral Hemorrhage/Stroke | LA, LV |
| Male | 61 | 23.2 | N/A | Head Trauma | LA, LV |
| Female | 53 | 41.7 | 52 | Anoxia/Drug Intoxication | LA, LV |

| | | | | | |
|---|---|---|---|---|---|
| Female | 60 | 24.2 | 60-65 | Cerebrovascular Accident/Intracerebral Hemorrhage/Stroke | LA, LV |
| Female | 64 | 21 | 60 | Cerebrovascular Accident/Intracerebral Hemorrhage/Stroke | LA, LV |

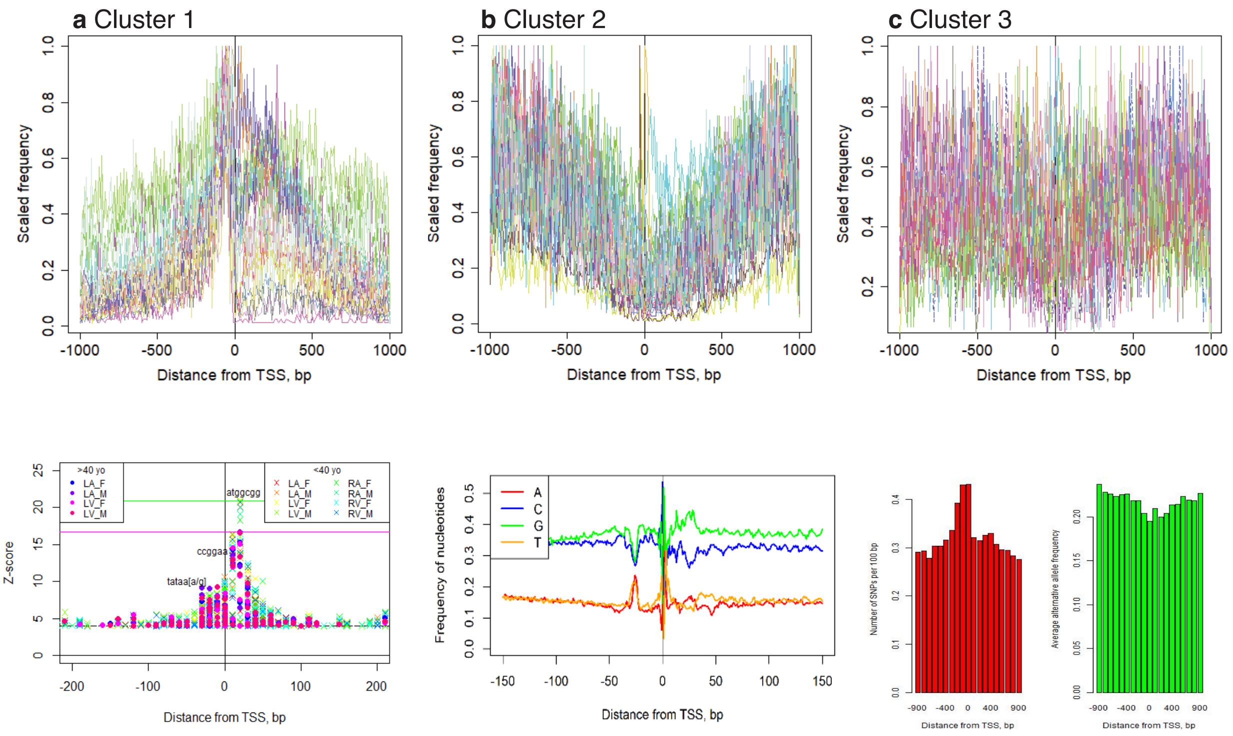


**Supplementary Fig. 1.** Position-specific frequencies of TFBS for three clusters: Cluster 1 (a), Cluster 2 (b), Cluster 3 (c). Motif frequencies are scaled (f/max(f)). (d) Z-scores of significant motifs (e) Nucleotide consensus around TSS (f)Number of SNPs and their average MAF in the vicinity of TSS.

Some of the transcription factors had a preferred binding position [5]. The resulting clusters are shown in **Supplementary Fig. 1**. **Supplementary Fig. 1a** (Cluster 1) contains TFBS with an affinity toward the core promoter area. Cluster 2 (**Supplementary Fig. 1b**) motifs had the opposite trend - they avoided the area. Cluster 3 (**Supplementary Fig. 1c**) did not have a positional binding preference. We have used public gene expression experiments to investigate the function of these clusters. We selected the genes with only Cluster 1, Cluster 2, or Cluster 3 binding sites in the promoter region. Generally, the presence of the Cluster 1 vs. Cluster 2 motifs resulted in lower average gene expression and higher variability of gene expression. The difference between gene expression in Clusters 1 and 2 ranges from 20% to 200% (for various gene expression sets). At the same time, the average expression of genes with Cluster 3 TF was 0.8-1.1 of Cluster 1. Therefore, Cluster 2 corresponds to the TF associated with broadly expressed genes, and Cluster 1 with more specific expression.

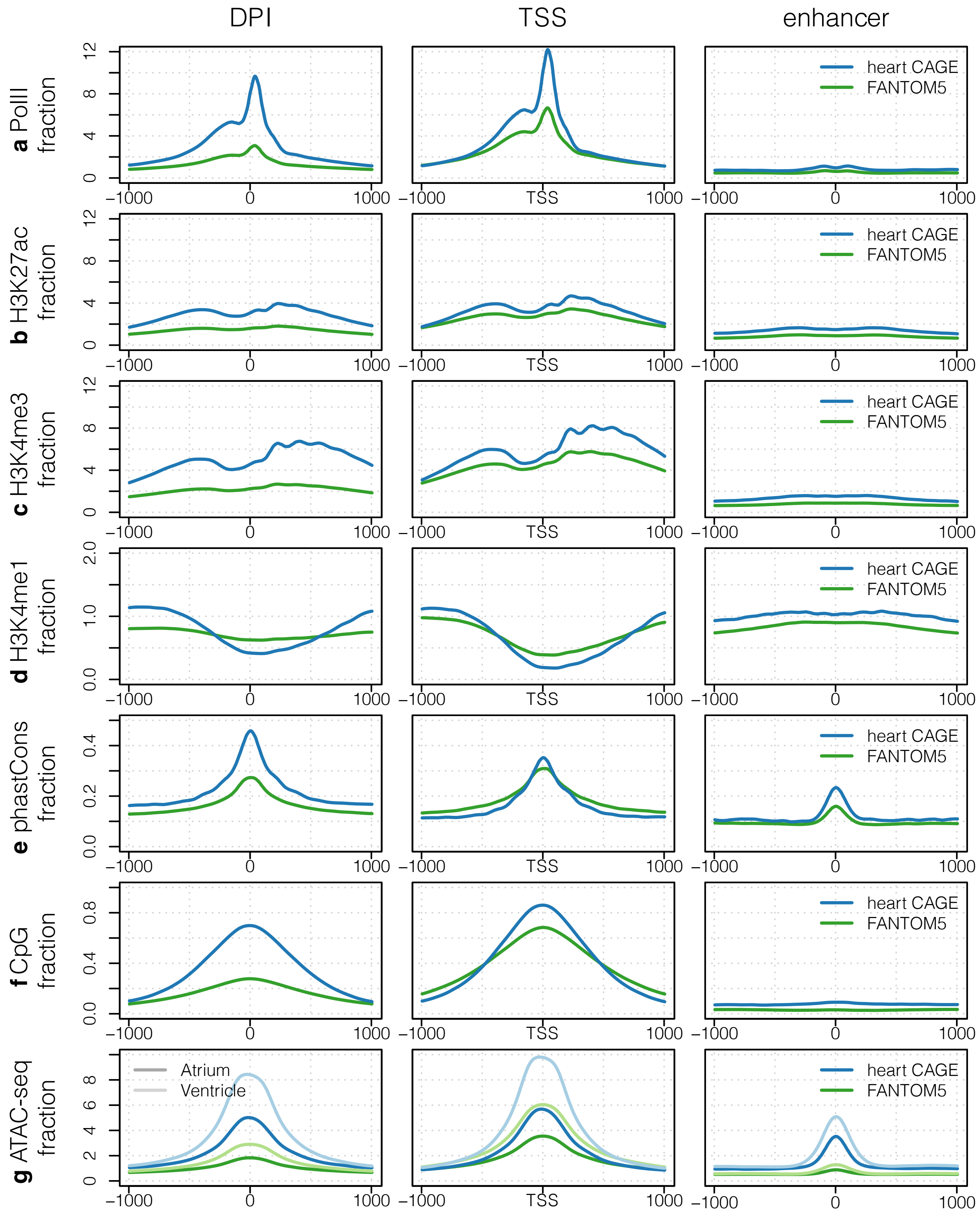


**Supplementary Fig. 2. Heart CAGE peaks have features of promoters and enhancers.** Signals and scores normalized by the number of clusters. **a-d,** Signal distribution of epigenetic marks based on ENCODE Chip-seq data. **e**, Conservation scores within defined regions, based on 100-way genomic alignment. **f**, CpG islands coverage of heart regulatory elements. In comparison to

FANTOM5 heart, DPI clusters have features of housekeeping CpG promoters. **g**, Heart ventricle and atrium ATAC-seq signal overlap with heart CTSS; atrium and ventricle ATAC-seq signal marked by dark and light colors, respectively.

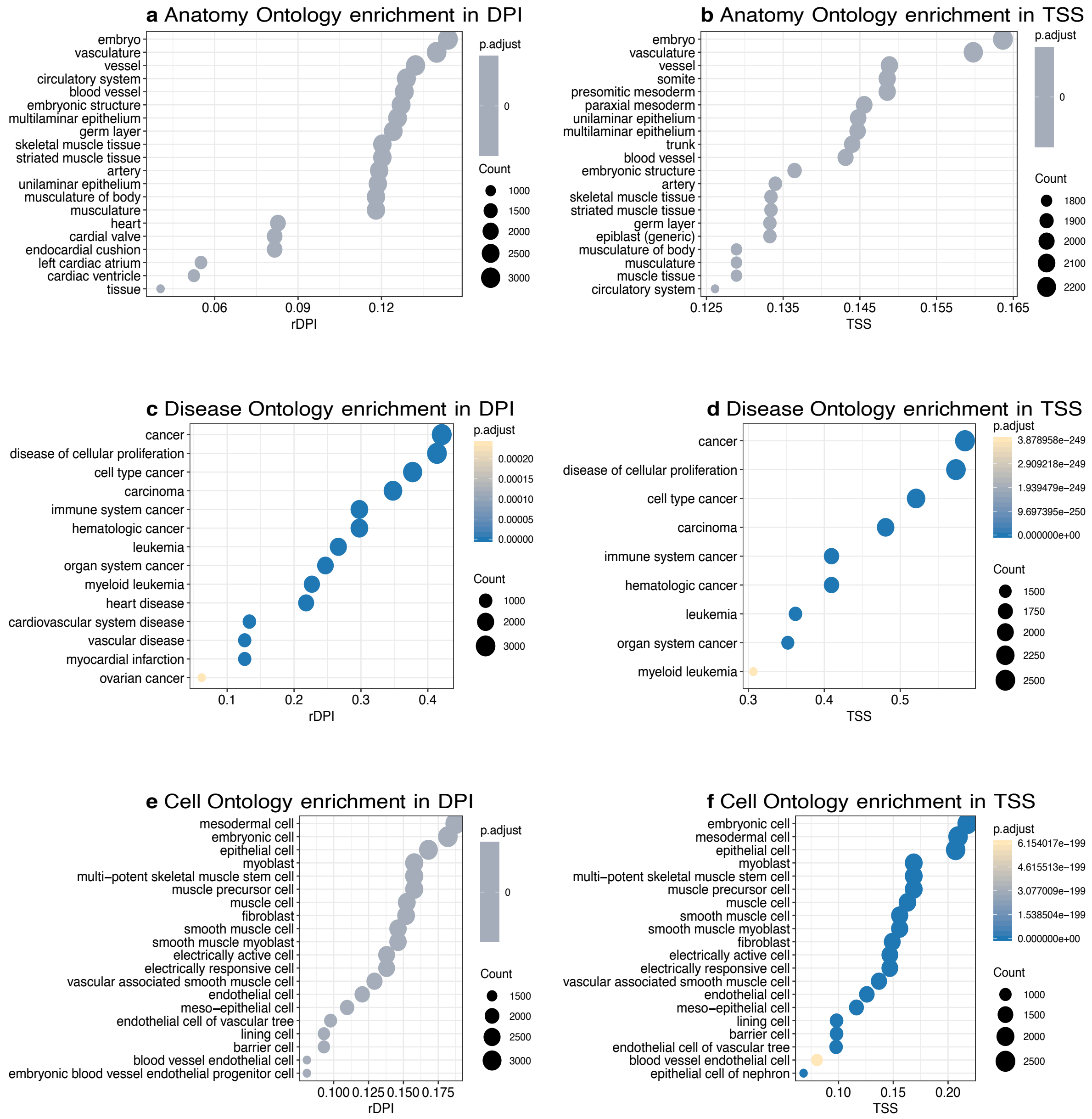


**Supplementary Fig. 3. Functional annotation of promoter activity defines TSS sub-clusters in the human heart.** Based on FANTOM5 ontologies intersection with heart CAGE. **a-b**, Uber Anatomy Ontology (UBERON) tags enrichment in heart CAGE peaks. Embryo specific expression profiles highly enriched. **c-d**, Disease Ontology enrichment for heart CAGE. **e-f**, Cell Type Ontology (CL) enrichment. These results are consistent and show that in the human heart there is a significant number of promoters with proliferation-related expression profiles.

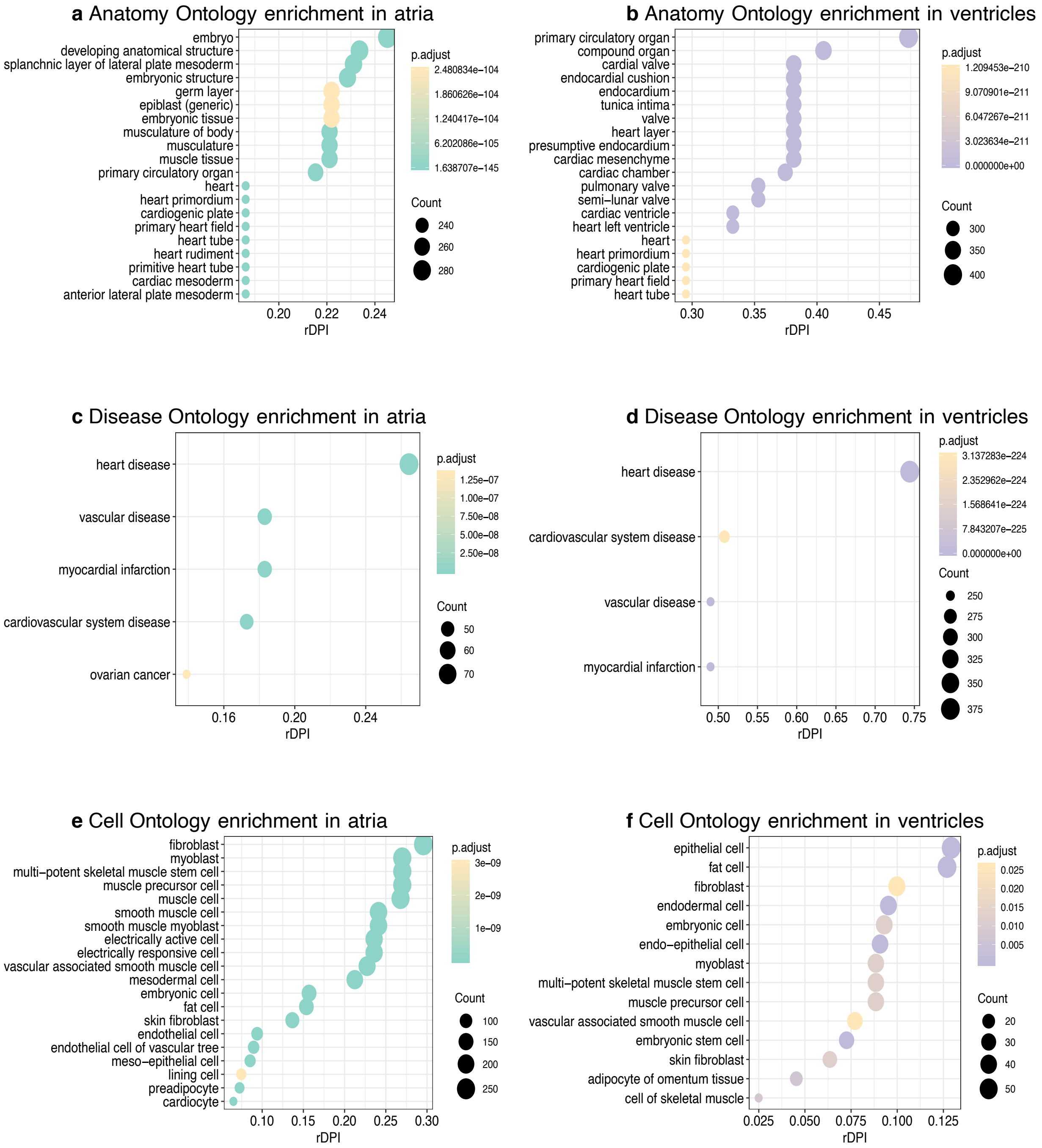


**Supplementary Fig. 4. Functional annotation of chamber-specific promoters based on FANTOM5 ontologies intersection with differentially expressed heart CAGE clusters. a-b**, UBERON; **c-d**, DOID; **e-f**, CL enrichment results. From this analysis, we found that embryo-type expression is atrium related, while ventricle expression profiles correlate mostly with heart samples from the FANTOM5 project.

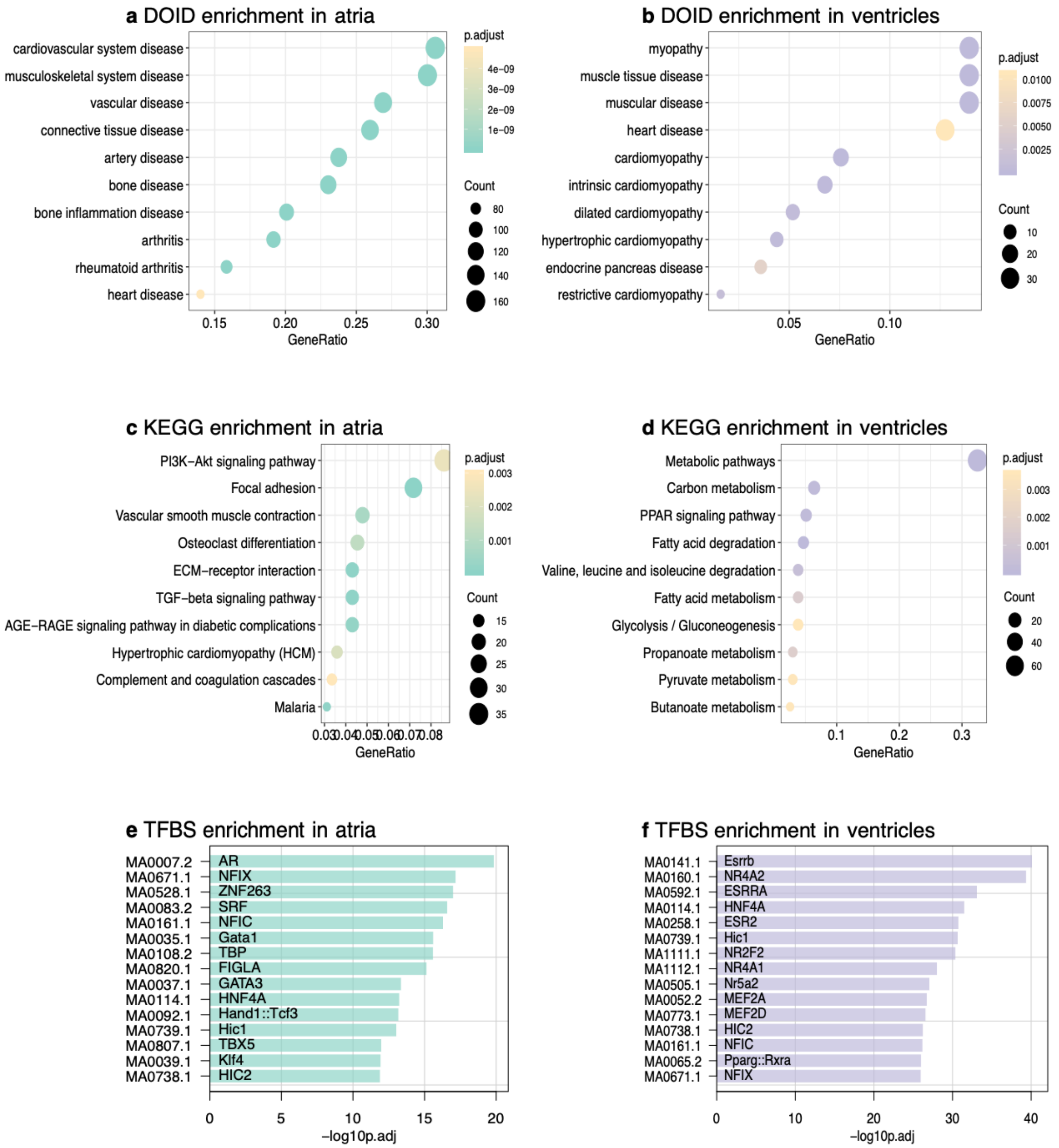


**Supplementary Fig. 5. Functional annotation and TF regulation of genes with chamber specific CAGE expression. a-b**, DOID enrichment results using gene ontology (https://disease-ontology.org/). **c-d**, KEGG pathways enrichment results. **e-f**, TF motifs enrichment in specific promoters.

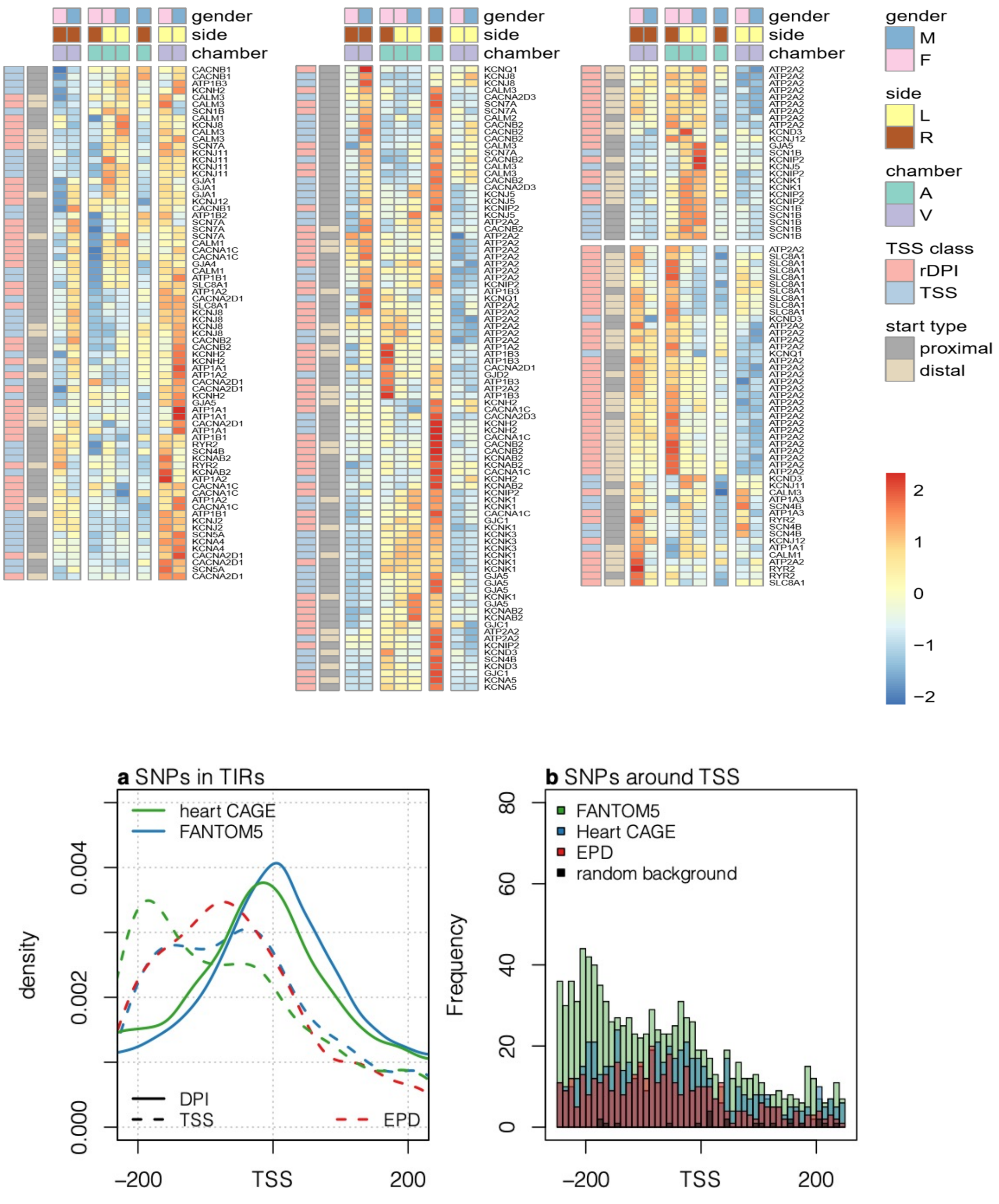


**Supplementary Fig. 6. Top. Promoter level expression profiles of electrophysiology genes.** We selected 40 genes related to cardiac electrophysiology and visualized its expression in samples younger than 40 years. Replicates were merged. The legend shows scaled TPM values. See the text for details. **Bottom. Heart GWAS SNPs overlap with CAGE. a**, position of SNP in transcription initiation regions (TIRs) defined by CAGE in comparison to EPD promoters. **b,** frequency of GWAS SNPs around TSS-like regions. Diseases related mutations have low frequency in heart promoters' regions and have a specific trend.

There is an interesting relationship between SNP density and TBFS of different clusters. Somewhat counterintuitively, the density of SNPs peaks in the promoter region is higher. However, this apparent excess of SNPs in promoters is misleading. The peak is achieved due to rare alleles (**Supplementary Fig. S1f**). Overall, there is a drop of sequence variability, if allele frequency is taken into account. Overall, functionally important regions are protected, since a mutation in a TBFS may be detrimental for binding affinity and negatively affect gene expression. Therefore, we analyzed the relationship between SNPs and TBFS of three clusters described above. We have focused on the region [TSS-300, TSS+100] and compared 4 datasets: allele frequencies of all SNPs in the area, and allele frequencies of SNPs that overlap with the positions of the TFBS of Clusters 1, 2, and 3. The average allele frequency of all SNPs is 0.208, for Cluster 1 it is 0.159, for Cluster 2 it is 0.212, and for Cluster 3 it is 0.190. To test the significance of those differences, we used the Kruskal-Wallis test, resulting in a p-value < 2.2e-16. Wilcoxon rank-sum test with continuity correction between Clusters 1 and 2 resulted in a p-value = 6.462e-11. Between Cluster 1 and all SNPs: p-value = 1.155e-10. Cluster 1 and all SNPs p-value = 8.166e-05. Therefore, we can conclude that motifs of Cluster 1 are not only frequent in the core promoter region, but they are also significantly depleted of the sequence variation.

### Differential expression analysis

Using the *glmFit* function from the *edgeR* package, we have identified differentially expressed genes between heart compartments, genders, and left and right sides of the heart. As expected, there were no significant differences between left and right (FDR=1). It is interesting to compare old and young datasets in terms of the maintenance of differences. We computed Pearson's correlation coefficients between logFC (log2 of the ratio of gene expression in ventricle to the atrium) and logCPM LogCPM (the log counts per million) between young and old adults. LogCPM is well preserved; the correlation is 0.94; LogFC is less correlated at 0.64.

**Supplementary Table 4. <40 y.o. ventricle to atrium expression**

| Gene | logFC | logCPM | LikelihoodRatio | PValue | FDR | ventricle/atrium |
|---|---|---|---|---|---|---|
| MYBPHL_109307003 | -8.4997 | 7.980678 | 42.3337 | 7.70E-11 | 8.52E-07 | 0.002763 |
| MYBPHL_109307024 | -5.81237 | 5.303411 | 41.21446 | 1.36E-10 | 8.52E-07 | 0.017795 |
| MYBPHL_109307035 | -6.21037 | 5.729147 | 40.94401 | 1.57E-10 | 8.52E-07 | 0.013505 |
| PRDM16_3069201 | 3.878869 | 4.885665 | 35.67417 | 2.33E-09 | 7.83E-06 | 14.71146 |
| SMYD2_214281224 | 4.635943 | 6.564233 | 35.61846 | 2.40E-09 | 7.83E-06 | 24.86325 |
| MASP1_187291700 | 2.967024 | 7.361751 | 34.64979 | 3.95E-09 | 9.54E-06 | 7.819216 |

| | | | | | | |
|---|---|---|---|---|---|---|
| LBH_30231535 | 3.656497 | 8.003915 | 34.579 | 4.09E-09 | 9.54E-06 | 12.61001 |
| ZNF638_71276600 | 2.786102 | 5.013287 | 34.2361 | 4.88E-09 | 9.95E-06 | 6.897636 |
| SMYD2_214281185 | 3.383738 | 5.343931 | 33.57154 | 6.87E-09 | 1.06E-05 | 10.43774 |
| HEY2_125749615 | 3.567777 | 5.504282 | 33.3297 | 7.78E-09 | 1.06E-05 | 11.8579 |
| RARRES2_150341620 | -2.94348 | 7.607074 | 33.16347 | 8.47E-09 | 1.06E-05 | 0.129995 |
| NMNAT2_183418350 | -3.95627 | 5.553327 | 33.15314 | 8.52E-09 | 1.06E-05 | 0.064423 |
| LMAN1L_74812844 | -3.14515 | 5.50495 | 32.92086 | 9.60E-09 | 1.06E-05 | 0.113036 |
| GALNT17_71131710 | 4.174761 | 5.698239 | 32.88053 | 9.80E-09 | 1.06E-05 | 18.06044 |
| GNAO1_56191487 | -2.87472 | 5.885506 | 32.84199 | 1.00E-08 | 1.06E-05 | 0.13634 |
| LMAN1L_74812834 | -2.86747 | 4.742627 | 32.76599 | 1.04E-08 | 1.06E-05 | 0.137027 |
| NMNAT2_183418380 | -3.44539 | 4.810515 | 32.48458 | 1.20E-08 | 1.10E-05 | 0.091798 |
| ZNF638_71276591 | 2.344729 | 5.841437 | 32.43713 | 1.23E-08 | 1.10E-05 | 5.07965 |
| HOOK2_12775527 | 2.418576 | 5.002573 | 32.34135 | 1.29E-08 | 1.10E-05 | 5.34643 |
| NDRG4_58464280 | 2.357284 | 7.964694 | 32.18087 | 1.40E-08 | 1.10E-05 | 5.124048 |

**Supplementary Table 5. >40 y.o. ventricle to atrium expression**

| Gene | logFC | logCPM | LR | PValue |
|---|---|---|---|---|
| SMYD2_214281224 | 5.364069 | 6.938725 | 23.21711 | 1.45E-06 |
| SMYD2_214281206 | 5.01836 | 6.075437 | 22.37627 | 2.24E-06 |
| LBH_30231535 | 3.660079 | 8.424985 | 21.71601 | 3.16E-06 |
| PITX2_110623076 | -3.87848 | 4.105782 | 21.59957 | 3.36E-06 |
| GALNT17_71131710 | 4.640096 | 5.15997 | 21.40853 | 3.71E-06 |

| | | | | |
|---|---|---|---|---|
| IRX5_54930995 | 4.489536 | 4.134616 | 20.77485 | 5.17E-06 |
| RHBDL3_32265830 | -3.22215 | 4.913746 | 20.35683 | 6.43E-06 |
| MYBPHL_109307035 | -4.31126 | 5.440804 | 20.08931 | 7.39E-06 |
| XPR1_180632086 | 3.384108 | 5.573022 | 20.0572 | 7.52E-06 |
| SMYD2_214281185 | 3.985877 | 4.582298 | 19.96597 | 7.88E-06 |
| NMNAT2_183418380 | -3.60492 | 4.357293 | 19.83731 | 8.43E-06 |
| GNAO1_56191487 | -2.99754 | 5.703087 | 19.77273 | 8.72E-06 |
| XPR1_180632057 | 3.193781 | 5.89745 | 19.67399 | 9.18E-06 |
| HAND1_154478226 | 3.563809 | 5.266911 | 19.63482 | 9.37E-06 |
| MYBPHL_109307024 | -3.95428 | 5.159488 | 19.58195 | 9.64E-06 |

### De-novo discovery of regulatory elements in CAGE defined promoters

We have restricted our dataset to the TSSs that are located upstream of the previously annotated open reading frames corresponding to the biologically validated protein-coding genes. There are 7,246 genes with 16,323 CAGE-based TSS, making approximately 2.25 alternative TSS per gene. Using the 7,246 protein-coding genes expressed in the heart, we divided all human genes into two categories. “Group 1” consisted of genes that were not expressed in the heart, according to the CAGE experiment. “Group 2” genes were expressed in the heart (not exclusively). Using all experiments from the FANTOM5 database, we calculated summary statistics for gene expression for genes in the two groups. To assess gene expression variability, we calculated the mean and coefficient of variation (CV) (a ratio of the standard deviation to its mean value) of $log2(CPM)$. Group 2 of genes had an overall high expression across all experimental conditions (almost twice higher than the genes that are not expressed in the heart). In heart, expression of the Group 2 genes was even higher (2.93 times higher than their expression not in heart, and 5.5 times higher than the expression of genes not expressed in heart, Group 1). Variability of the Heart group was the smallest in the heart (0.98 vs. 2.62). Group 1 genes were more variably expressed than the Group 2 (6.59 vs. 2.62). Therefore, Group 1 genes could be described as tissue-specific constitutively expressed genes. Regulatory regions of the tissue-specific constitutively expressed genes were supposed to have a particular structure. To investigate promoter regions of these genes, we used *cis*Express [6,7] to find regulatory elements in promoters that were significantly associated with expression in different parts of the heart. cisExpress is based on two critical assumptions: (i) the function of promoter motifs is position-specific, and (ii) gene expression data provide reasonable measurements of transcript abundance and reflect promoter activity.

The most significant motifs in the "Young" (<40 y.o.) and in "Old" (>40 y.o.) datasets were: "atggcgg" at +10 and +20 [8] and "tataa[a/g]" -30 and -20, "ccggaa" and "ggaagt" at -10 (Fig. S1d). The relationship between sequence and motif was more significant in the Young dataset. The identified motifs have been well studied across multiple organisms [8,9]. "The Ying and Yang" motif (YY1) "atggcgg", located around position 20 downstream from the TSS, is usually present in the housekeeping genes [8,10], which is also consistent with the classification of TFBS presented above and low variability of the gene expression of the selected genes. Motif "ccggaa" is a binding site of the E-twenty-six (ETS) transcription factor family; these TFs have a demonstrated relationship with cardiovascular health [11], explicitly playing a role in the development of abdominal aortic aneurysms, they are expressed in the embryonic heart and regulate distal Gata4 cardiac enhancer, functioning as a master regulator of the development of the cardiovascular system [12]. The ETS transcription factors play an essential role in pathophysiological angiogenesis and endothelial cell reprogramming [13,14]. TATA-box, a binding site for TBP (the TATA-binding protein), is located approximately 20 nt upstream from the TSS (Fig. S1e).

### Biological significance

Kalirin RhoGEF kinase (KALRN) is located on human chromosome 3. The KALRN gene encodes several isoforms by using alternative transcription start and termination sites.[15,16] CAGE sequencing analysis confirmed the heart activity of at least two of these alternative starts of transcription, 552,218 bp apart. The two start sites are located in positions 124,032,456 (Site A) and 124,584,674 (Site V); Based on CAGE, Site A is approximately equally active in the atrium and ventricle (15.66:12.15), while Site V is almost 3-fold more active in the ventricle (8.94:25.13). We have assessed the cell-type-specific expression for isoforms A and V from the FANTOM5 database.

**Supplementary Table 6. FANTOM5 data on cell-type specificity of the expression of isoforms "A" and "V" in heart tissues.**

| **Experiment** | **"A"** | **"V"** |
|---|---|---|
| Heart mitral valve adult | 7.773160 | 0.896903 |
| Heart pulmonic valve adult | 15.949997 | 4.706556 |
| Heart tricuspid valve adult | 4.995850 | 3.330567 |
| Heart adult diseased post-infarction | 3.877075 | 5.305470 |

| | | |
|---|---|---|
| Heart adult diseased | 5.109989 | 7.664984 |
| Heart adult pool | 1.609841 | 8.049207 |
| Heart fetal pool | 6.05812 | 12.93768 |

**Supplementary Table 7. Expression of isoforms of KALRN in cell lines.**

| CELL | Isoform | |
|---|---|---|
| | "A" | "V" |
| medial temporal gyrus | 56.64032 | 1.937231 |
| occipital cortex | 53.99163 | 4.302783 |
| pineal gland | 47.48883 | 0.836447 |
| retina | 7.951958 | 60.73308 |
| parietal lobe | 24.79278 | 2.41658 |
| medial frontal gyrus | 26.29388 | 4.337429 |
| optic nerve | 32.01677 | 0 |
| hippocampus | 17.179 | 2.753119 |
| eye - muscle medial | 26.78787 | 55.06396 |
| temporal lobe | 16.31552 | 0.626879 |
| spinal cord | 14.96951 | 2.893579 |

| | | |
|---|---|---|
| brain | 17.05135 | 2.427592 |
| cerebellum | 10.27876 | 0.332605 |
| eye - muscle inferior rectus | 26.5603 | 45.87689 |
| olfactory region | 19.86941 | 1.146312 |
| frontal lobe | 19.79686 | 1.125888 |
| postcentral gyrus | 20.48623 | 2.179386 |
| insula | 19.21059 | 1.798035 |
| stomach | 0 | 17.08757 |
| occipital pole | 17.6605 | 1.419147 |
| Urethra | 16.18584 | 0 |
| globus pallidus | 10.39909 | 2.87002 |
| paracentral gyrus | 16.84005 | 2.471095 |
| locus coeruleus | 10.01832 | 3.132035 |
| colon carcinoma | 0 | 9.701225 |
| ARPE-19 EMT induced with TGF-beta and TNF-alpha | 1.732247 | 0.022641 |
| amniotic membrane cells | 0 | 7.52201 |
| substantia nigra | 7.766123 | 1.375549 |
| amygdala | 8.936237 | 0.202767 |

| | | |
|---|---|---|
| pancreas | 0 | 12.22831 |
| corpus callosum | 15.47872 | 3.316869 |
| MCF7 breast cancer cell line response to EGF | 0 | 1.659163 |
| heart - pulmonic valve | 15.95 | 4.706556 |
| merkel cell carcinoma | 7.930918 | 0 |
| thalamus | 11.11592 | 6.109915 |
| aorta | 11.17958 | 0 |
| parietal cortex | 11.74411 | 0.734007 |
| occipital lobe | 8.060284 | 0.524798 |
| skeletal muscle - soleus muscle | 17.02404 | 6.384015 |
| pituitary gland | 4.164821 | 10.02819 |
| cruciate ligament | 9.725643 | 0 |
| MCF7 breast cancer cell line response to HRG | 0 | 1.396966 |
| nucleus accumbens | 11.71739 | 2.905912 |
| pons | 9.553477 | 0.774606 |
| heart | 4.163756 | 8.489335 |
| small intestine | 0.783189 | 6.772362 |
| umbilical cord | 8.346005 | 0 |

| | | |
|---|---|---|
| medulla oblongata | 12.03097 | 7.860616 |
| achilles tendon | 8.266116 | 0 |
| neuroepithelioma | 0 | 8.26496 |
| salivary gland | 8.579284 | 0.571952 |
| cerebral meninges | 10.08558 | 2.269255 |
| tongue | 1.870315 | 7.297162 |
| penis | 9.668019 | 2.148449 |
| adrenal gland | 7.275337 | 0 |
| hepatocellular carcinoma | 0 | 4.172943 |
| K562 erythroblastic leukemia response to hemin | 0 | 0.976016 |
| HES3-GFP Embryonic Stem cells c cardiomyocytic induction | 0.220012 | 1.330301 |
| heart - mitral valve | 7.77316 | 0.896903 |
| placenta | 0 | 6.450166 |
| diaphragm | 0.802563 | 7.223066 |
| tongue epidermis | 9.238847 | 3.079616 |
| small cell lung carcinoma | 3.034198 | 0 |
| small cell gastrointestinal carcinoma | 5.35849 | 11.34739 |
| rectum | 0 | 5.563969 |

| | | |
|---|---|---|
| Neurons | 3.283584 | 0.073388 |
| medulloblastoma | 3.874709 | 0.061136 |
| H9 Embryoid body cells c melanocytic induction | 0.20793 | 0.96096 |
| Rinderpest infection | 0.051012 | 1.476599 |
| carcinoid | 0.994877 | 3.813321 |
| Mast cell | 0.050028 | 1.626028 |
| signet ring carcinoma | 0 | 3.334558 |
| retinoblastoma | 4.676441 | 0 |
| eye - muscle superior | 19.49044 | 24.07643 |
| eye | 1.271171 | 5.847388 |
| ductus deferens | 4.485059 | 0 |
| Intestinal epithelial cells | 0 | 4.19476 |
| mature adipocyte | 2.127476 | 0.051882 |
| prostate | 4.3532 | 0.362767 |
| immature langerhans cells | 0.121073 | 2.916069 |
| liver | 0.324123 | 3.047509 |
| testis | 2.985438 | 0.279229 |
| adipose tissue | 3.722861 | 0 |

| | | |
|---|---|---|
| hIPS | 2.574758 | 0.479096 |
| smooth muscle | 3.61837 | 0 |
| tonsil | 0.601054 | 4.207375 |
| amygdala - adult | 9.486379 | 5.928987 |
| spleen | 1.470697 | 3.938116 |
| xeroderma pigentosum b | 0 | 3.471479 |
| Clontech Human Universal Reference Total RNA | 3.943126 | 0.550204 |
| bone marrow | 4.048956 | 0.674826 |
| uterus | 3.412652 | 1.109171 |
| vagina | 3.230994 | 0 |
| epididymis | 3.558287 | 0.418622 |
| Myoblast differentiation to myotubes | 0.428722 | 0.003588 |
| parotid gland | 3.083319 | 0 |
| diencephalon | 4.319791 | 1.43993 |
| submaxillary gland | 2.848404 | 0 |
| thyroid | 4.220778 | 6.173836 |
| seminal vesicle | 2.645357 | 0 |
| Fingernail including nail plate eponychium and hyponychium | 2.614592 | 5.229184 |

| | | |
|---|---|---|
| cervical cancer | 0 | 1.822882 |
| colon | 0.391678 | 1.880054 |
| Adipocyte | 0.73307 | 0.179982 |
| Chondrocyte | 0.022756 | 1.155976 |
| breast | 2.525425 | 0 |
| caudate nucleus | 5.475117 | 4.214661 |
| Prostate Epithelial Cells | 0 | 1.428368 |
| rectal cancer | 0 | 2.32748 |
| cervix | 3.464676 | 1.154892 |
| Saos-2 osteosarcoma treated with ascorbic acid and BGP to induce calcification | 0.173897 | 0.486093 |
| skin | 1.893682 | 0.315614 |
| cerebrospinal fluid | 7.308843 | 5.11619 |
| left atrium | 3.813956 | 1.634553 |
| Astrocyte | 0.871078 | 0.020469 |
| Lymphatic Endothelial cells response to VEGFC | 0.303049 | 0.009205 |
| adipose | 1.232446 | 0.216688 |
| pancreatic carcinoma | 0 | 1.956747 |

| | | |
|---|---|---|
| osteosarcoma | 0 | 1.378384 |
| appendix | 2.328708 | 0.388118 |
| breast carcinoma | 0.312057 | 1.593155 |
| neuroectodermal tumor | 0.424422 | 1.468304 |
| trachea | 0.945135 | 2.205316 |
| esophagus | 0.743246 | 2.442093 |
| mesenchymal stem cells | 0.24447 | 0.014989 |
| heart - tricuspid valve | 4.99585 | 3.330567 |
| chronic myelogenous leukemia | 0 | 0.72902 |
| mesodermal tumor | 1.547428 | 0 |
| vein | 1.753587 | 0.219198 |
| left ventricle | 1.516963 | 3.033927 |
| hIPS bCCl2 | 0.304842 | 1.163632 |
| eye - vitreous humor | 1.457391 | 0 |
| CD34 cells | 0 | 0.591659 |
| chorionic membrane cells | 0.311278 | 1.139678 |
| Hepatocyte | 0.81256 | 1.619628 |
| artery | 6.793445 | 5.434756 |

| | | |
|---|---|---|
| SABiosciences XpressRef Human Universal Total RNA | 0.668179 | 1.870901 |
| squamous cell carcinoma | 0.815389 | 0.13031 |
| lung | 1.226988 | 0.556274 |
| Neural stem cells | 1.870171 | 1.08882 |
| gastric adenocarcinoma | 0 | 0.773664 |
| duodenum | 0 | 0.735649 |
| Olfactory epithelial | 0.511856 | 0 |
| neuroblastoma | 0.611179 | 0.104103 |
| Preadipocyte - perirenal | 0.984268 | 0 |
| pleomorphic hepatocellular carcinoma | 0.933836 | 0 |
| hepatoma | 0 | 0.889726 |
| bladder | 1.412335 | 0.543206 |
| malignant trichilemmal cyst | 0 | 0.849495 |
| H1 embryonic stem cells differentiation to CD34 b HSC | 0.247238 | 0.529768 |
| skeletal muscle | 1.821818 | 2.398746 |
| gall bladder carcinoma | 0 | 0.548619 |
| Fibroblast - skin normal | 0.385345 | 0 |
| Universal RNA - Human Normal | 0.738145 | 1.476291 |

| Tissues Biochain | | |
|---|---|---|
| adenocarcinoma | 0 | 0.706499 |
| Neutrophils | 0 | 0.407228 |
| lung adenocarcinoma | 0 | 0.483098 |
| Melanocyte - dark | 0.388973 | 0 |
| small cell cervical cancer | 0.667046 | 0 |
| Skeletal Muscle Satellite Cells | 0.429961 | 0.047563 |
| dura mater | 3.799014 | 3.138316 |
| Osteoblast - differentiated | 0.381038 | 0 |
| Whole blood 8ribopure 9 | 0 | 0.229783 |
| glioblastoma | 0.409985 | 0.035184 |
| mucinous adenocarcinoma | 0 | 0.635808 |
| Endothelial Cells - Microvascular | 0.335356 | 0 |
| Fibroblast - skin spinal muscular atrophy | 0.250009 | 0 |
| choriocarcinoma | 0.207127 | 0.525028 |
| throat | 1.995349 | 2.384617 |
| carcinosarcoma | 0.532356 | 0 |
| iPS differentiation to neuron | 0.320759 | 0.396653 |

| | | |
|---|---|---|
| Preadipocyte - omental | 0.299798 | 0 |
| gall bladder | 2.172891 | 1.690027 |
| Hep-2 cells treated with Streptococci | 0 | 0.19324 |
| CD19 b B Cells | 0 | 0.14261 |
| Bronchial Epithelial Cell | 0 | 0.178615 |
| embryonic kidney | 0.071647 | 0.404756 |
| Corneal Epithelial Cells | 0 | 0.255631 |
| ovary | 0.698812 | 0.262055 |
| putamen | 3.964971 | 4.159671 |
| bile duct carcinoma | 0 | 0.300683 |
| tubular adenocarcinoma | 0 | 0.423252 |
| mesenchymal stem | 0.421014 | 0 |
| Renal Epithelial Cells | 0.237107 | 0 |
| thymic carcinoma | 0.397464 | 0 |
| Myoblast | 0.225323 | 0 |
| Fibroblast - Villous Mesenchymal | 0.222734 | 0 |
| prostate cancer | 0 | 0.272252 |
| Preadipocyte - breast | 0.247561 | 0.056287 |

| | | |
|---|---|---|
| salivary acinar cells | 0 | 0.220684 |
| acute myeloid leukemia | 0 | 0.078866 |
| Dendritic Cells - plasmacytoid | 0 | 0.217623 |
| alveolar cell carcinoma | 0 | 0.3573 |
| leukemia | 0 | 0.354174 |
| Saos-2 osteosarcoma cell line | 0.183457 | 0.376808 |
| Osteoblast | 0.124389 | 0.308022 |
| testicular germ cell embryonal carcinoma | 0 | 0.154213 |
| kidney | 1.043108 | 0.833644 |
| renal cell carcinoma | 0.207105 | 0 |
| glioma | 0.585755 | 0.292878 |
| Endothelial Cells - Lymphatic | 0.167852 | 0 |
| Placental Epithelial Cells | 0 | 0.159444 |
| Fibroblast - Gingival | 0.076084 | 0 |
| Retinal Pigment Epithelial Cells | 0 | 0.135882 |
| somatostatinoma | 0.268021 | 0 |
| gamma delta positive T cells | 0 | 0.186746 |
| normal intestinal epithelial | 0.262764 | 0 |

| | | |
|---|---|---|
| Nucleus Pulposus Cell | 0.148113 | 0 |
| nasal epithelial cells | 0 | 0.147824 |
| Gingival epithelial cells c donor3 8GEA15 9.CNhs11903.11379-118B2.hg38.nobarcode | 0 | 0.255513 |
| Meningeal Cells | 0.041353 | 0.188069 |
| glassy cell carcinoma | 0 | 0.249418 |
| Skeletal muscle cells differentiated into Myotubes | 0.186286 | 0.045866 |
| mesenchymal precursor cell - ovarian cancer right ovary | 0.108644 | 0 |
| Prostate Stromal Cells | 0.139034 | 0 |
| maxillary sinus tumor | 0.238259 | 0 |
| mucinous cystadenocarcinoma | 0 | 0.230358 |
| Fibroblast - Aortic Adventitial | 0.086449 | 0 |
| CD133 b stem cells | 0 | 0.159887 |
| Fibroblast - skin c normal | 0.15682 | 0 |
| thymus | 0.155857 | 0 |
| Smooth Muscle Cells - Bronchial | 0.152216 | 0 |
| leiomyoma | 0.11975 | 0 |
| Smooth Muscle Cells - Uterine | 0.146289 | 0 |

| | | |
|---|---|---|
| B lymphoblastoid | 0 | 0.118218 |
| Keratocytes | 0.11461 | 0 |
| Lens Epithelial Cells | 0.182605 | 0.07208 |
| Smooth Muscle Cells - Brain Vascular | 0.10853 | 0 |
| Gingival epithelial cells | 0 | 0.13012 |
| ductal cell carcinoma | 0 | 0.12822 |
| bronchial squamous cell carcinoma | 0 | 0.103073 |
| CD4 b T Cells | 0 | 0.101664 |
| Smooth Muscle Cells - Pulmonary Artery | 0 | 0.100515 |
| non T non B acute lymphoblastic leukemia | 0.16912 | 0 |
| Mammary Epithelial Cell | 0 | 0.096517 |
| Papillotubular adenocarcinoma | 0 | 0.165582 |
| Clear cell carcinoma | 0.114291 | 0 |
| Fibroblast - Lymphatic | 0.093292 | 0 |
| Renal Mesangial Cells | 0.09247 | 0 |
| Epidermoid carcinoma | 0 | 0.111159 |
| Renal Proximal Tubular Epithelial Cell | 0 | 0.089474 |
| embryonic pancreas | 0 | 0.075678 |

| | | |
|---|---|---|
| Peripheral Blood Mononuclear Cells | 0 | 0.086277 |
| Ciliary Epithelial Cells | 0.215513 | 0.300507 |
| Fibroblast - Periodontal Ligament | 0.044998 | 0.102985 |
| serous adenocarcinoma | 0 | 0.080975 |
| Melanocyte - light | 0 | 0.078357 |
| serous cystadenocarcinoma | 0 | 0.134321 |
| lung adenocarcinoma | 0 | 0.131816 |
| Perineurial Cells | 0.092011 | 0 |
| Pericytes | 0.074178 | 0 |
| CD14 b Monocytes | 0.034124 | 0.01385 |
| Fibroblast - Conjunctival | 0.084802 | 0 |
| H9 Embryonic Stem cells | 0.119553 | 0.051359 |
| Urothelial Cells | 0 | 0.057301 |
| Schwann Cells | 0.06269 | 0 |
| teratocarcinoma | 0.159695 | 0.102765 |
| Preadipocyte - subcutaneous | 0.056762 | 0 |
| large cell lung carcinoma | 0 | 0.069121 |
| Synoviocyte | 0 | 0.055875 |

| | | |
|---|---|---|
| Smooth Muscle Cells - Umbilical Artery | 0.047008 | 0 |
| Endothelial Cells - Vein | 0.052336 | 0 |
| Smooth Muscle Cells - Umbilical Vein | 0.046581 | 0 |
| Fibroblast - Choroid Plexus | 0.044664 | 0 |
| Fibroblast - Dermal | 0.027071 | 0 |
| Mesenchymal Stem Cells | 0 | 0.01458 |
| Skeletal Muscle Cells | 0.11416 | 0.13934 |
| Smooth Muscle Cells - Internal Thoracic Artery | 0.033547 | 0 |
| Keratinocyte - epidermal | 0.031552 | 0 |
| Preadipocyte - visceral | 0.031049 | 0 |
| Renal Cortical Epithelial Cells | 0 | 0.034656 |
| CD4 bCD25 bCD45RA- memory regulatory T cells | 0 | 0.010302 |
| Tracheal Epithelial Cells | 0.023895 | 0 |
| mesothelioma | 0.009744 | 0 |
| Aortic smooth muscle cell response to IL1b | 0.006881 | 0 |
| Aortic smooth muscle cell response to FGF2 | 0.014977 | 0.017955 |
| Smooth Muscle Cells - Tracheal | 0.027769 | 0.027769 |

| | | |
|---|---|---|
| Iris Pigment Epithelial Cells | 0.137207 | 0.137207 |
| pagetoid sarcoma | 0.18332 | 0.18332 |
| peripheral neuroectodermal tumor | 0.185936 | 0.185936 |
| mesenchymal precursor cell - bone marrow | 0.231359 | 0.231359 |
| Skin - palm | 8.058326 | 8.058326 |
| eye - muscle lateral | 14.72943 | 14.72943 |
