## Supplementary figures and images for "Human library of cardiac promoters and enhancers"

### Supplement 4 Fig. 6

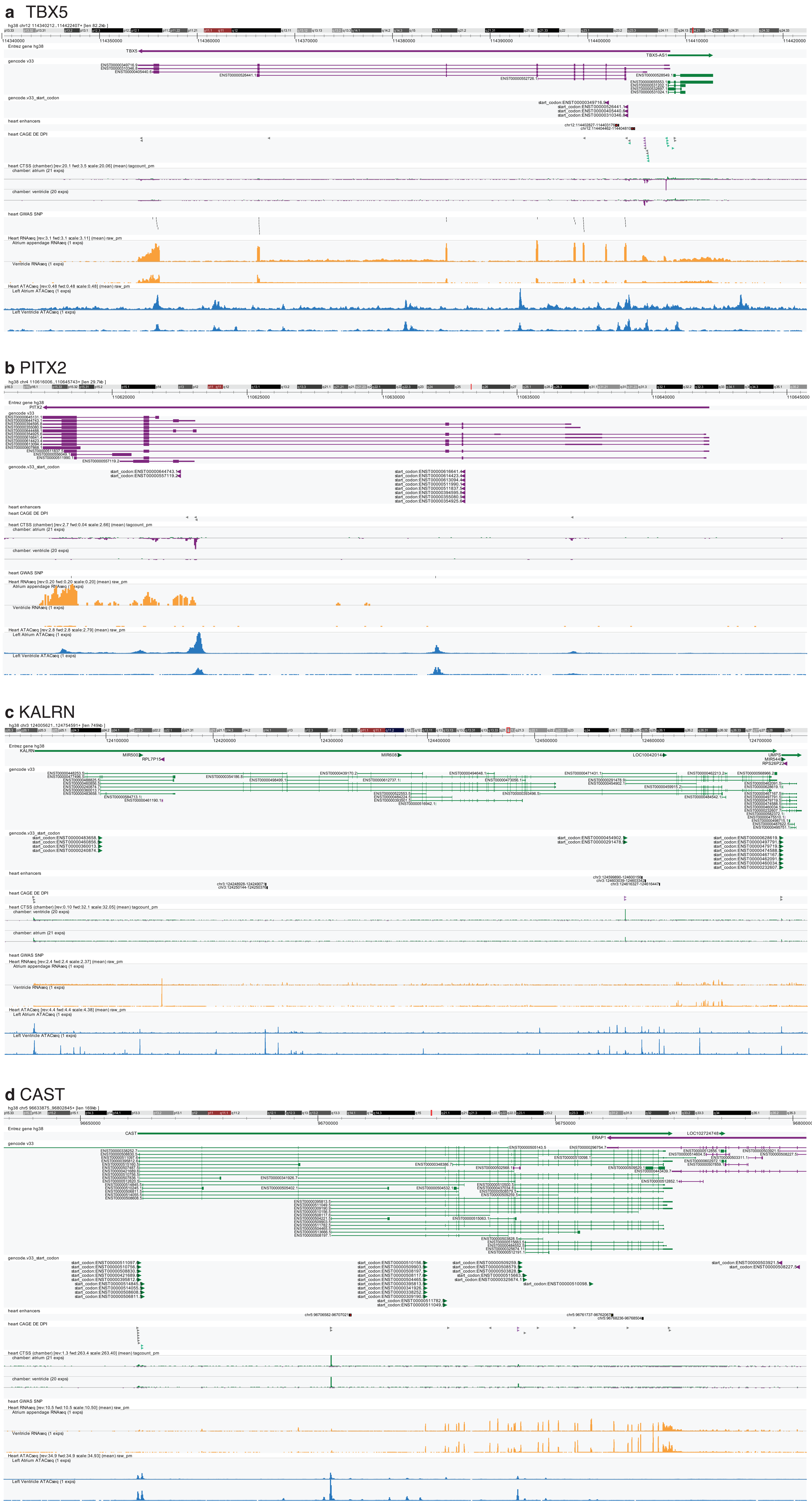

**e** IRF6

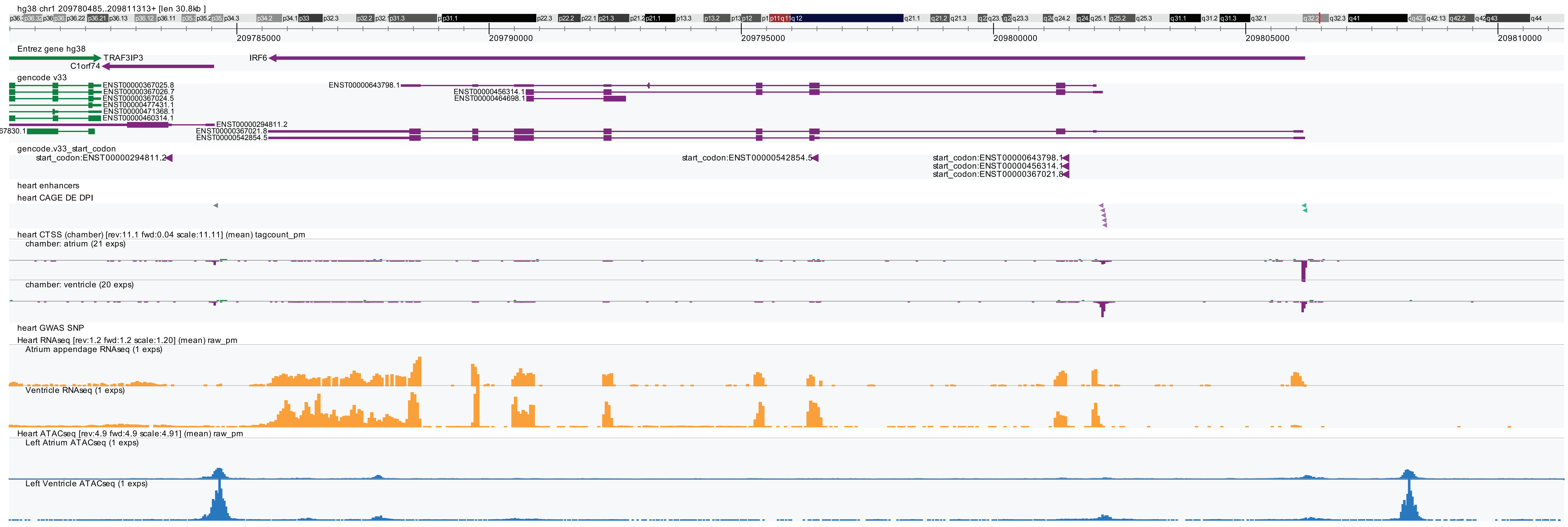


**f** ABLIM1

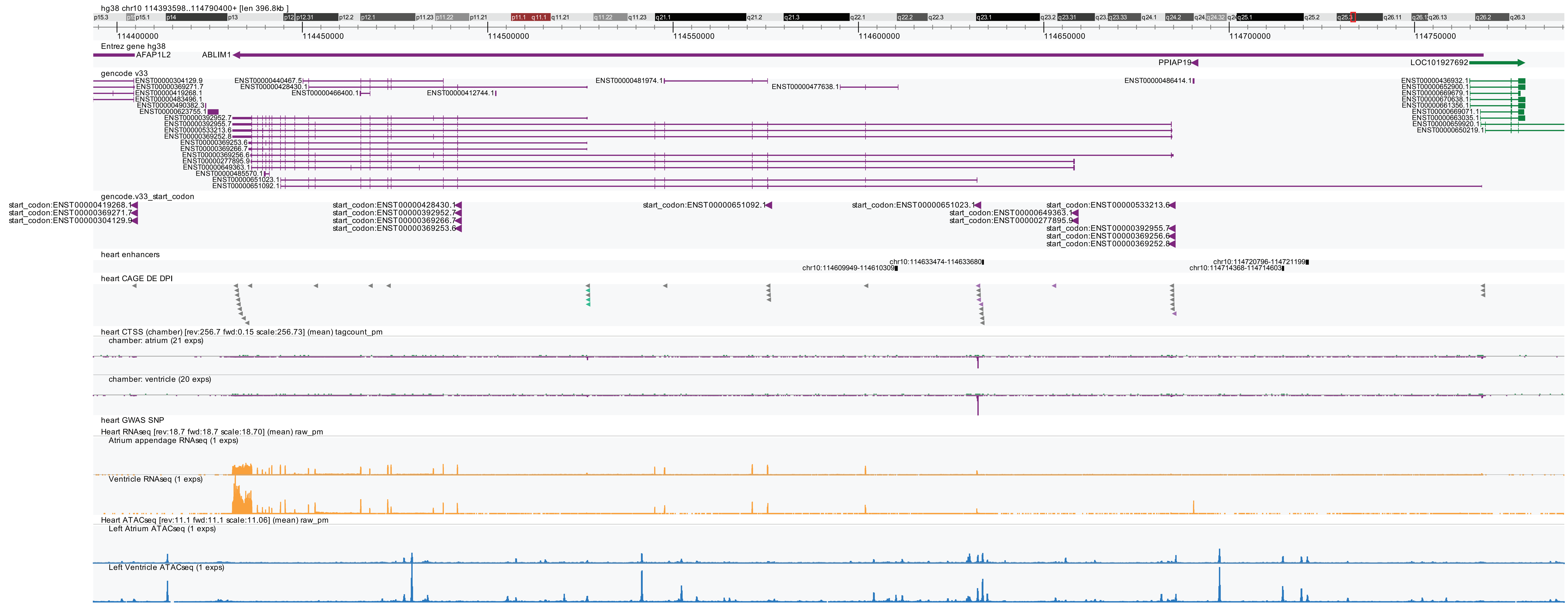


**g** AMOTL1

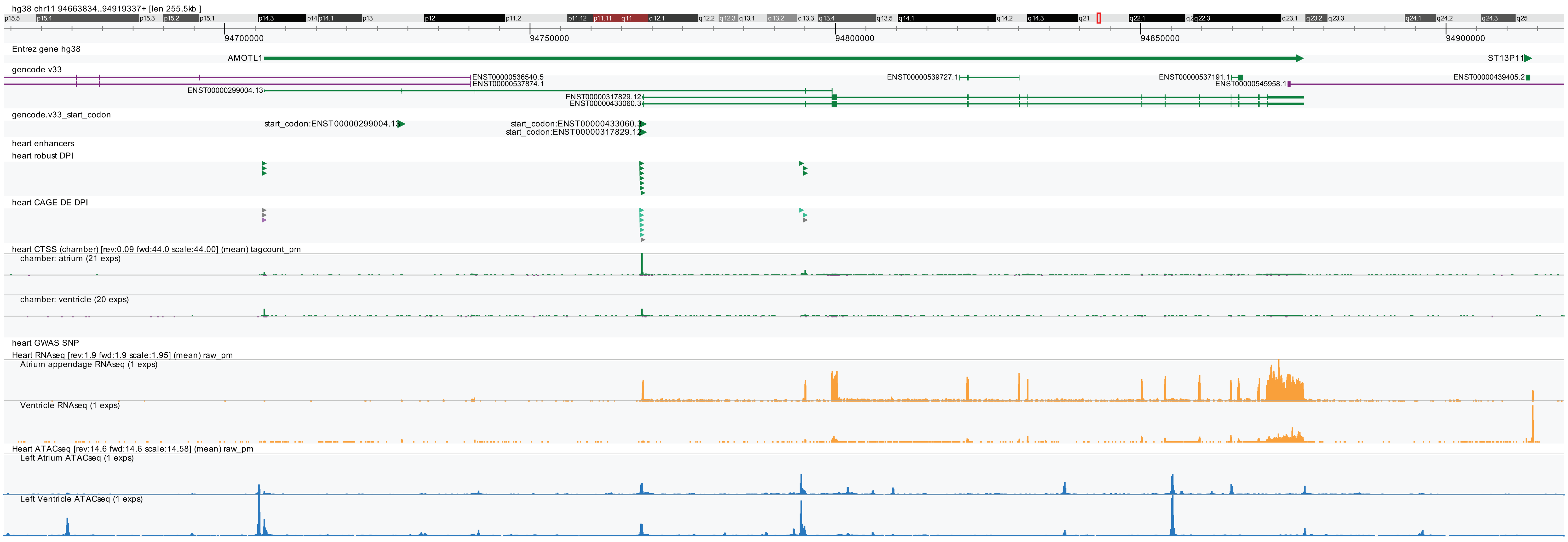


**h** RGS3

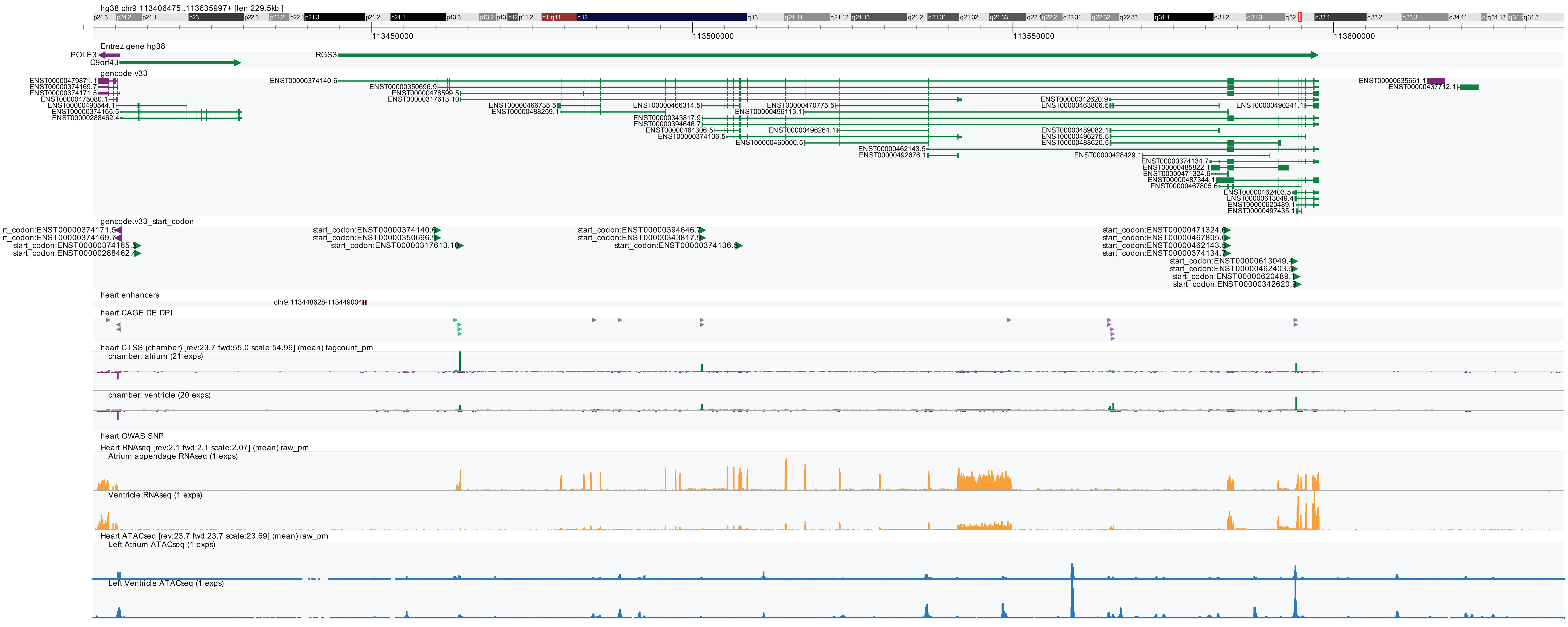
