## Supplement 5 Zenbu User Manual for "Human library of cardiac promoters and enhancers"

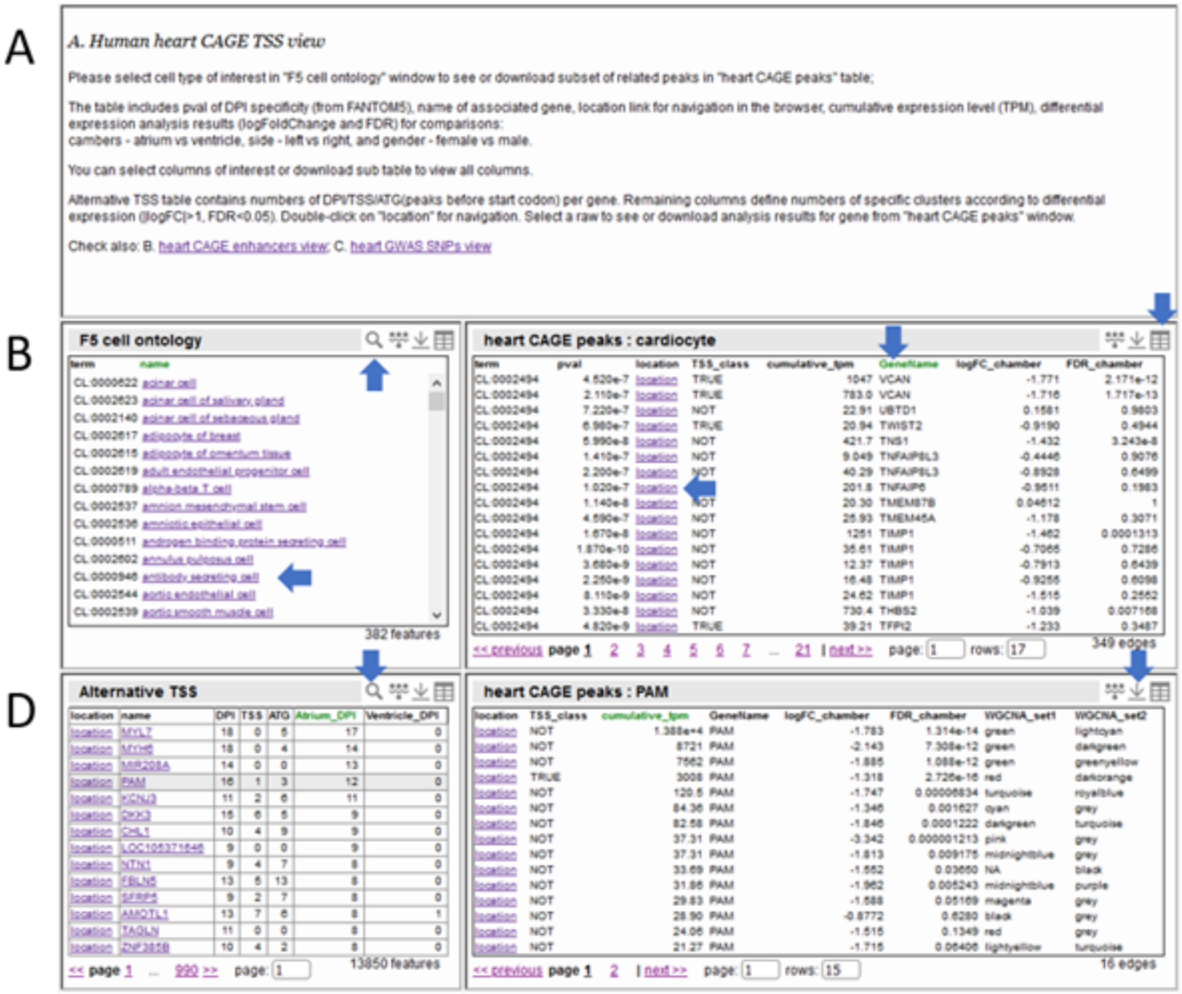


**Human heart CAGE TSS view on Zenbu reports page.** A - brief description and links to other Heart library pages. B - FANTOM5 cell ontology table based on CAGE peaks overlap between FANTOM5 and heart CAGE. Select cell type of interest to see related heart CAGE peaks in table C. C - heart CAGE peaks annotation table. Includes differential expression, co-expression, gene name data. You can add or remove columns, sort by value, and select location of the feature for navigation in the browser (tough click). D - Gene annotation table by heart CAGE clusters. You can search genes of interest or order by columns (additional columns are available). This table helps to find cases of alternative TSS usage in heart. By clicking on the row of interest you can get information about CAGE clusters related to the gene in table E. You can also download these tables.

https://fantom.gsc.riken.jp/zenbu/reports/#Human_Heart_CAGE_A

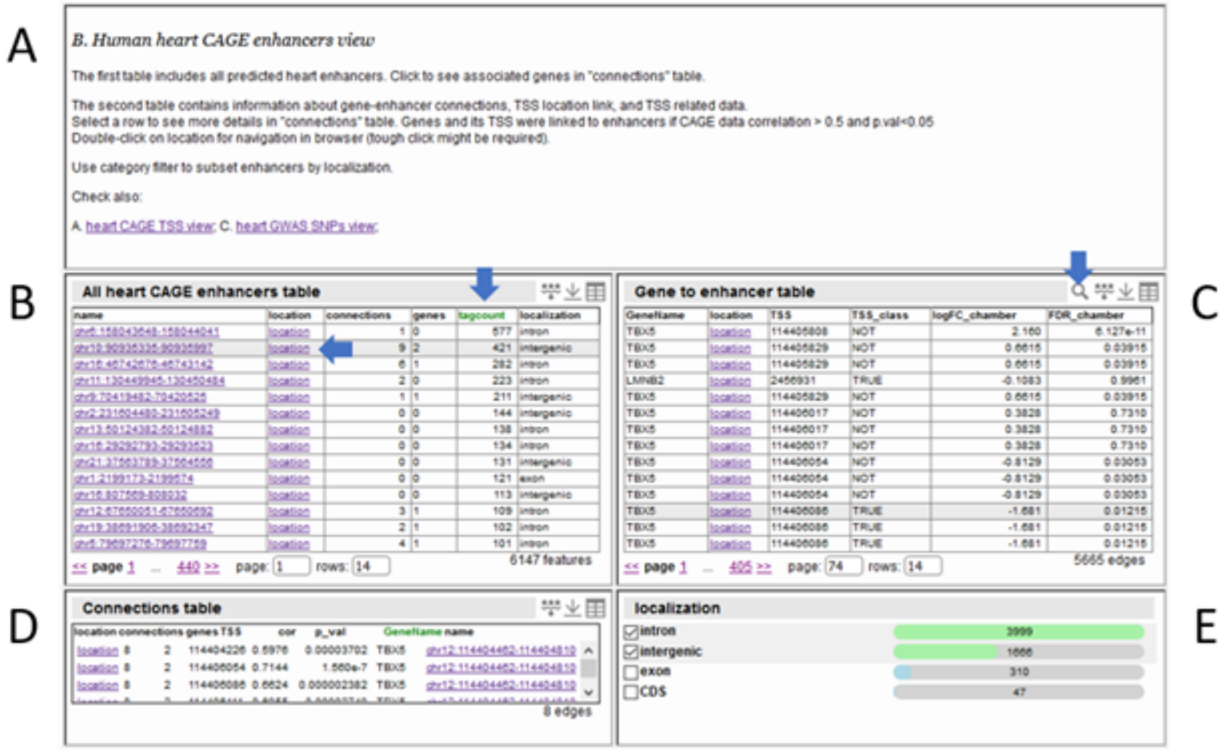


**Human heart CAGE enhancer view on Zenbu reports page.** A - brief description and links to other Heart library pages. B - heart CAGE enhancer table, includes all predicted regions, numbers of mapped CAGE tag counts, numbers of enhancer-to-DPI, and enhancer-to-gene connections based on expression correlation and distance, location link to enhancer (tough click). C - Gene annotation table by heart CAGE enhancers. If the gene was connected to the heart CAGE enhancer it will be available in this table. Location link to connected DPI. E - category filter for enhancer localization for table C. For example, keep genes annotated by only intronic and intergenic enhancers. Select a row of interest in table B or C to see correlation test results in table D (location link for enhancer).

https://fantom.gsc.riken.jp/zenbu/reports/#Human_Heart_CAGE_B

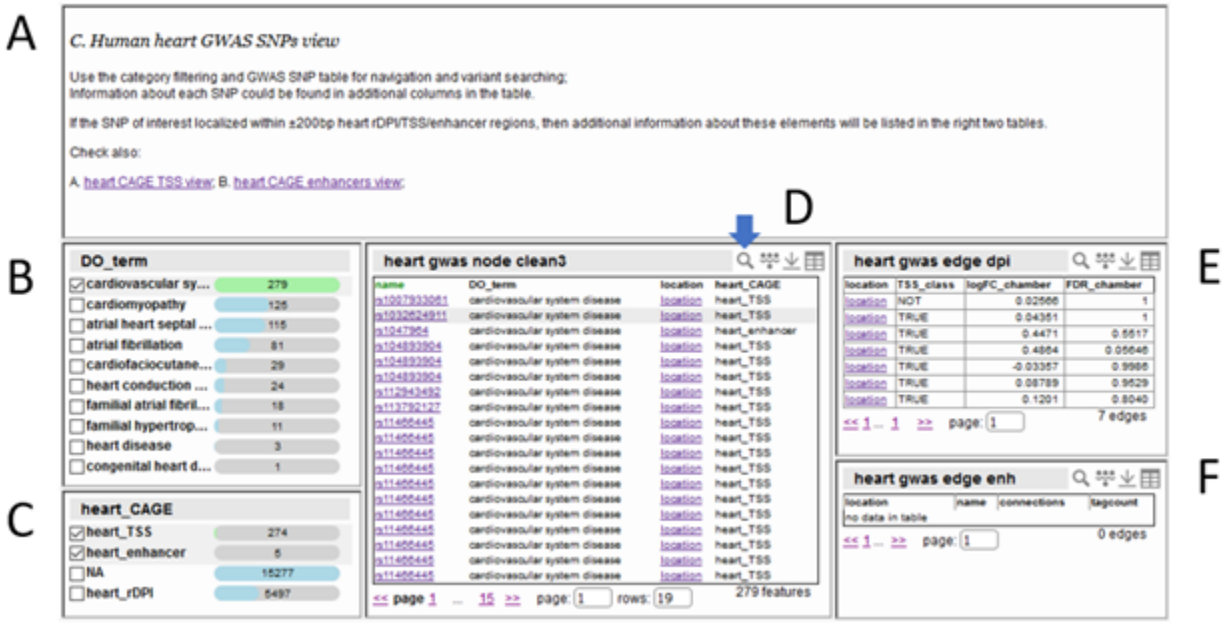

**Human heart GWAS SNPs view on the Zenbu reports page.** A - brief description and links to other Heart library pages. B and C - category filters for SNP table D. In table D additional columns are available, including study title, authors, year, PMC, PubMed number. Find SNP of interest and select a row to see information about overlapped heart CAGE promoters and enhancers in tables E and F, respectively.

https://fantom.gsc.riken.jp/zenbu/reports/#Human_Heart_CAGE_C

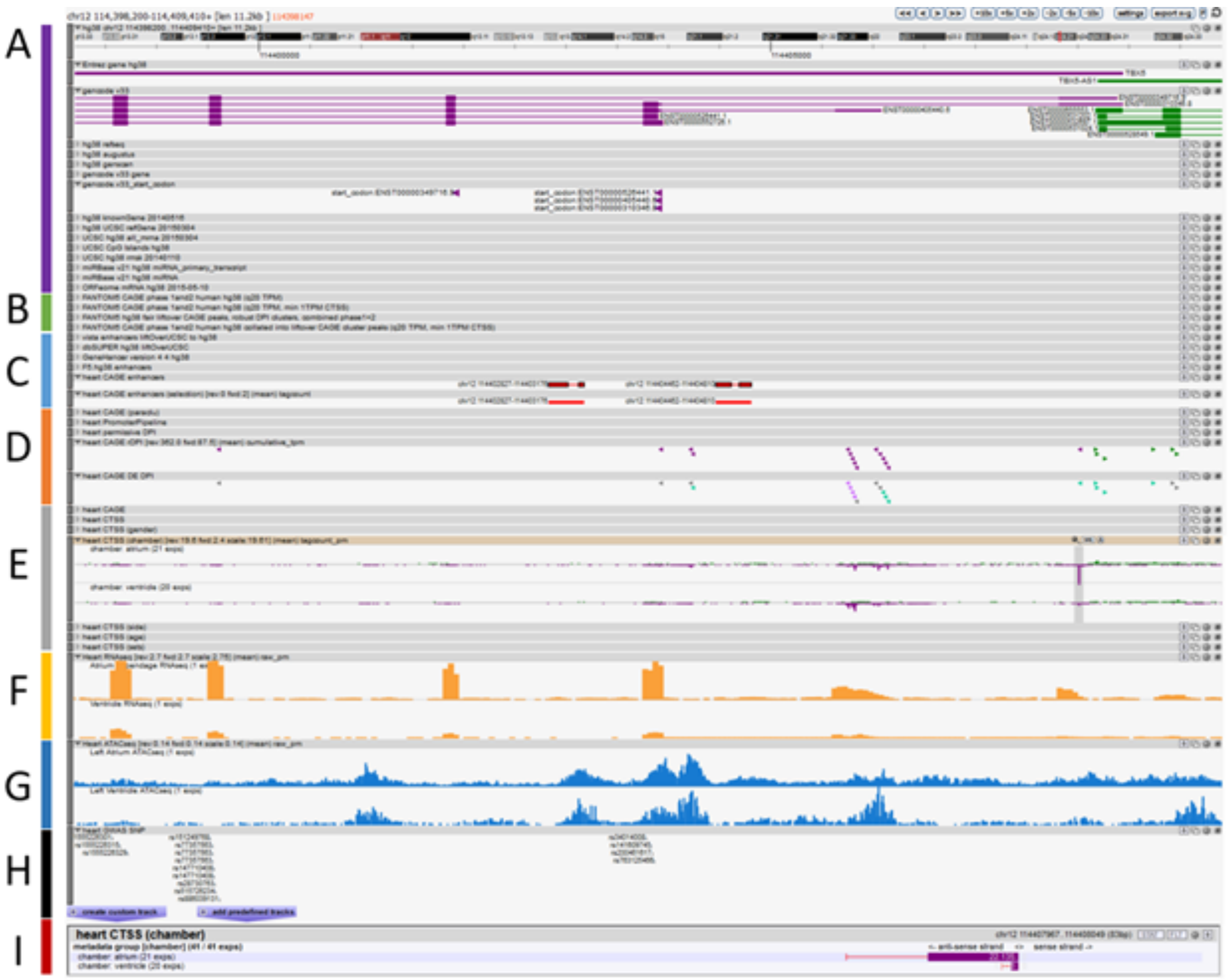


Zenbu browser view on Heart CAGE library of promoters and enhancers. A - genome and gene model tracks. B - FANTOM5 CAGE data tracks and DPI peaks. C - enhancer annotation tracks from multiple sources, including heart CAGE, ‘selection’ track will highlight selected enhancer on the enhancer view page. D - heart CAGE peaks called by DPI, Promoter Pipeline, paraclu algorithms. Heart CAGE rDPI track will highlight selected robust DPI clusters on the TSS view page. Heart CAGE DE DPI track use grey, green, and purple color for stable, atrium and ventricle specific robust DPI (|logFC|>1, FDR<0.05), respectively. E - grouped heart CAGE experiments by chamber (ventricle and atrium), by side (left, right), by age, by gender, by experimental set. Heart CAGE track allows quality filtration of mapped reads. F - RNAseq experiments for atrial appendage and ventricle from GEO GSE128188 and GSE116250 (non-failing), CPM normalized. G - ATACseq tracks from Broad Institute’s Cardiovascular Disease Knowledge Portal liftovered on hg38. H - heart GWAS SNP track. Selected SNP will be highlighted on the SNP view page. I - expression barplot for the selected track (for example, heart CAGE for chambers in E).

https://fantom.gsc.riken.jp/zenbu/gLyphs/#config=mRP7PgjCNhKi9gn0ajOwvC
